# CrGEF1, CrGAP1, and CrGDI2 function as key regulators of ROP-GTPase mediated modulation of alkaloid biosynthesis in *Catharanthus roseus*

**DOI:** 10.1101/2024.12.02.625791

**Authors:** Anuj Sharma, Sruthi Mohan, Priyanka Gupta, Durgesh Parihar, Dinesh A. Nagegowda

**Affiliations:** Molecular Plant Biology and Biotechnology Lab, CSIR-Central Institute of Medicinal and Aromatic Plants, Research Centre, Bengaluru-560065; Academy of Scientific and Innovative Research (AcSIR), Ghaziabad-201002, India

**Keywords:** ROP-GTPase regulatory proteins, Guanine nucleotide exchange factors, GTPase activating proteins, Guanine nucleotide dissociation inhibitors, Rho of plant, monoterpene indole alkaloids, *Catharanthus roseus*

## Abstract

ROP-GTPase Regulatory Proteins (RGRPs) have been shown to control plant morphogenesis, development and immunity, however, their role in specialized metabolism is hitherto not known. Here, we demonstrate that specific RGRPs control monoterpene indole alkaloid (MIA) biosynthesis by interacting with distinct Rho of Plants (ROP) in *Catharanthus roseus*. Among the five Guanine nucleotide Exchange Factors (GEFs), four GTPase-activating proteins (GAPs), and two GDP dissociation inhibitors (GDIs) identified in the *C. roseus* genome, only CrGEF1, CrGAP1, and CrGDI2 specifically interacted with CrROP3 and CrROP5. These RGRPs displayed distinct cytosolic and/or membrane localization patterns, with their transcripts predominantly expressed in aerial tissues. Functional studies revealed that *CrGEF1* acts as a positive regulator of MIA biosynthesis, as its silencing led to a reduction in MIA production, while overexpression enhanced MIA levels. Conversely, CrGAP1 and CrGDI2 function as negative regulators, with their silencing resulting in increased MIA production and their overexpression causing reduced MIA levels. Notably, terminal truncated forms of these RGRPs showed interaction with CrROP3 or CrROP5 but failed to influence MIA biosynthesis, underscoring the importance of these domains in their regulatory functions. Overall, our findings uncover a previously unexplored mechanism by which distinct RGRPs coordinate with specific ROPs to regulate transcription factors and fine-tune MIA biosynthesis in *C. roseus*.

**Significance:** Plants fine-tune their metabolic pathways to adapt to environmental and physiological cues, balancing primary and specialized metabolism. Here, we uncover a feedback regulatory mechanism by which ROP-GTPase regulatory proteins (RGRPs) control alkaloid biosynthesis in *Catharanthus roseus*. We found that CrGEF1, CrGAP1, and CrGDI2 specifically interact with ROP-GTPases CrROP3 and CrROP5, modulating the monoterpene indole alkaloid (MIA) pathway. This RGRP-ROP module plays a crucial role in dynamically regulating MIA levels, where CrGEF1 promotes MIA production while CrGAP1 and CrGDI2 act as negative regulators. These findings highlight the importance of RGRP-ROP circuitry in control of alkaloid biosynthesis in *C. roseus*, and specialized metabolism in plants.

## INTRODUCTION

Plants produce a wide array of natural products, often referred to as Plant Specialized Metabolites (PSMs), which play crucial roles in their survival and fitness, particularly under challenging environmental conditions, and also facilitate communication with surrounding organisms. The biosynthesis of these structurally diverse PSMs is an intricately regulated process that requires the coordinated involvement of numerous signaling pathways and regulatory elements (1). *Catharanthus roseus,* commonly known as Madagascar periwinkle, is a renowned medicinal plant belonging to the Apocynaceae family. It produces over 130 monoterpene indole alkaloids (MIA) (2), including the well-known anticancer bisindole alkaloids vincristine and vinblastine, the antihypertensive ajmalicine, and the sedative serpentine (3, 4). Besides being a rich source of pharmaceutically significant PSMs, *C. roseus* has been a great model for investigating alkaloid biosynthesis and its regulation. The formation of MIA in *C. roseus* leaves is a highly complex process, involving more than 50 biosynthetic steps. These steps are distributed across various cell layers of the leaf and involve a diverse array of components, including pathway enzymes, transcription factors (TFs), intra- and intercellular signaling molecules, and transporters (5). The precise coordination of signaling pathways and regulatory factors is crucial for this intricate process.

It has been shown that the TFs involved in regulation of MIA biosynthesis are in turn affected by protein geranylgeranylation, a post-translational modification involving the attachment of an isoprene unit to a protein (6, 7). This modification influences protein functionality by affecting subcellular localization, interactions with other proteins, and enzymatic activity (8, 9). Inhibition of protein geranylgeranylation has been shown to negatively impact the expression of TFs and pathway genes, resulting in reduced MIA accumulation in *C. roseus*, indicating the involvement of geranylgeranylated proteins (7, 10). Among the prenylated proteins, the Rho of plant (ROP) family of small GTPases, as *bona fide* signalling molecules, modulate essential cellular processes in plants, including growth, development, and responses to environmental stimuli (11–13). However, the exact role of ROPs in PSM biosynthesis was not well-studied. Our recent study clearly demonstrated that specific ROPs, CrROP3 and CrROP5, which contain the C-terminal geranylgeranylation motif “CSIL,” interact with protein geranylgeranyl transferase (PGGT-I, undergo geranylgeranylation, and positively regulate the expression of TFs and pathway genes thus enhancing MIA biosynthesis in *C. roseus* (6). Given the central role of CrROP3 and CrROP5 in controlling *C. roseus* MIA biosynthesis, knowing the precise spatiotemporal regulation of their activity is crucial to manifest the modulation of MIA accumulation as per the need of the stimuli. Hence, understanding how activation and inactivation of specific ROPs (in this case CrROP3 and CrROP5) is controlled is essential for unravelling the signaling circuitry involved in MIA modulation in *C. roseus*, and PSM biosynthesis in plants.

Studies have revealed that ROP signaling in plants is tightly regulated by accessory proteins like ROP-GTPase regulatory proteins (RGRPs), which include ROP-specific guanine nucleotide exchange factors (GEFs), GTPase-activating proteins (GAPs), and guanine nucleotide dissociation inhibitors (GDIs). These regulatory proteins modulate the cycling of ROPs between their active, GTP-bound, and inactive, GDP-bound states, thereby fine-tuning ROP-mediated signaling. ROP-GEFs activate ROP signaling by facilitating the exchange of GDP for GTP (12, 14). ROP-GEFs possess PRONE (Plant-specific ROP Nucleotide Exchanger) domains, which are unique to plant GEFs and significantly enhance the nucleotide exchange rate, promoting ROP activation over a thousand-fold (15). Though generally viewed as activators, ROP-GEFs may also perform nucleotide exchange-independent roles, revealing complexity in plant signaling (16). Unlike ROP-GEFs, ROP-GAPs regulate ROP signaling by facilitating GTP hydrolysis, controlling signal timing and strength. They contain a GAP domain and a CRIB motif essential for their function (17). Whereas ROP-GDIs regulate ROPs by controlling their membrane localization and sequestration, signaling balance, and cellular stability (18). ROP-GDIs contain a RhoGDI domain that binds Rho-GTPases, inhibiting activation and aiding in cytosolic recycling (19). While the role of ROP and RGRP circuitry in various primary processes is widely studied in different plants(12, 14, 20), its involvement in spatiotemporal activation and inactivation in the context of secondary metabolism remain unexplored.

In this study, we present, for the first time, a regulatory feedback loop that controls the biosynthesis of MIA in *C. roseus*. This loop involves the interconnected ROP proteins CrROP3 and CrROP5, and RGRPs CrGEF1, CrGAP1, and CrGDI2. While CrGEF1 positively influences MIA biosynthesis, CrGAP1 and CrGDI2 negatively regulate it. These specific RGRPs individually interact with CrROP3 or CrROP5, and regulate the expression of key TFs and pathway genes, thereby controlling MIA biosynthesis.

## RESULTS

### *Catharanthus roseus* Genome Encodes for 11 ROP-GTPase Regulatory Proteins (RGRPs)

To comprehensively identify RGRPs within the *C. roseus* genome, a two-pronged approach was employed. Firstly, a query search was conducted using the Medicinal Plant Genomics Resource (MPGR) database (http://medicinalplantgenomics.msu.edu/4058.shtml). This was followed by BLAST (BLASTp) search against the assembled *C. roseus* genome (21, 22) using characterized RhoGEFs, RhoGAPs, and RhoGDIs sequences from other plant species. To ensure high-confidence identifications, transcripts with low abundance Fragments Per Kilobase Million (FPKM), short length, or incomplete open reading frames (ORFs) were excluded. This stringent filtering resulted in 11 full-length transcripts, which encode five GEF, four GAP, and two GDI proteins exhibiting high similarity to known plant RGRPs. The genes encoding CrGEFs, CrGAPs, and CrGDIs exhibit complex gene structures with multiple introns and are unevenly distributed across five of the eight chromosomes (Fig. 1A-C) (SI Appendix, Fig. S1). Further, sequence analysis of CrGEFs revealed the presence of a conserved PRONE domain composed of three subdomains flanked by hyper-variable N- and C-terminal regions, which are known to regulate protein-protein interactions and subcellular localization(Fig. 1D) (SI Appendix, Fig. S2) (15). Similarly, CrGAPs possess conserved CRIB and GAP domains with variable terminal regions that facilitate functional diversity through differential regulation and interactions with signaling partners (Fig. 1E) (SI Appendix, Fig. S3) (17). Whereas CrGDIs contain the characteristic RhoGDI domain with variability in their terminal regions, which have been shown to fine-tune GDI’s regulatory role by modulating Rho protein interactions (23). (Fig. 1F) (SI Appendix, Fig. S4).

**Figure 1.**
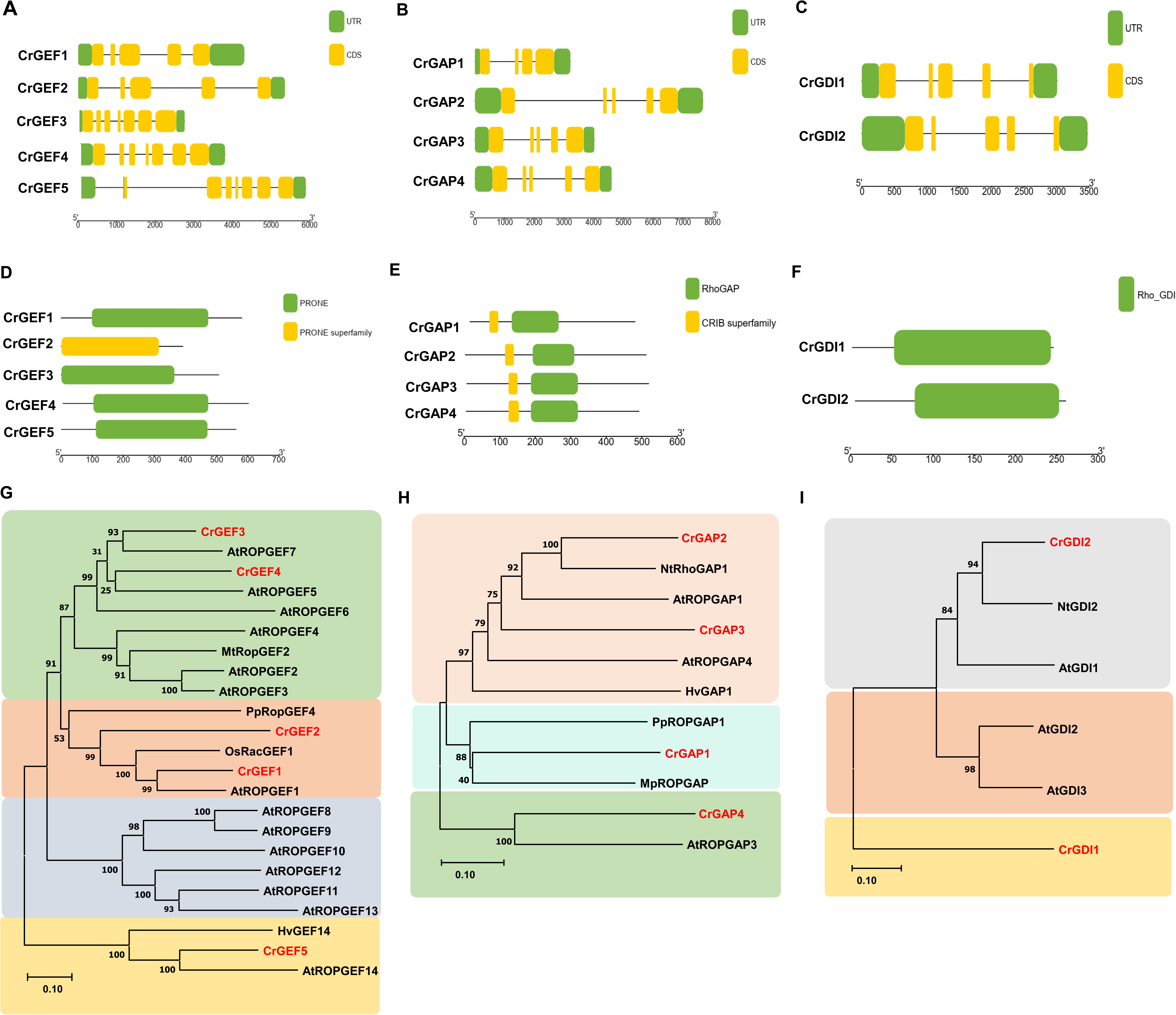
Gene Structure, Conserved Motif, and Phylogenetic Analysis of *Catharanthus roseus* RGRPs. (A-C) Schematic gene structure of *CrGEFs, CrGAPs* and *CrGDIs* depicting exons (in yellow) and introns (black line) along with untranslated regions (UTR, in green). The gene structure was made using TB tools (Chen et al. 2020). (D-F) Conserved domain structure of CrGEFs, CrGAPs and CrGDIs. Amino acid sequences of CrGEFs, CrGAPs and CrGDIs was analysed using NCBI conserved domain search (https://www.ncbi.nlm.nih.gov/Structure/cdd/wrpsb.cgi). The domain structure was made using TB tools (Chen et al. 2020). (G-I) Phylogenetic relationship of CrGEFs, CrGAPs and CrGDIs with GEFs, GAPs and GDIs of others plants. The tree was constructed using neighbour-joining method with bootstrap value of 1,000 runs using MEGA 11 software. Accession number of sequences used to construct the tree are provided in (SI Appendix, Table S1).

Phylogenetic analysis of CrGEFs with characterized GEFs from other plants, categorized them into four distinct groups (Fig. 1G). CrGEF1 and CrGEF2 were placed in Group 2 and displayed high sequence similarity to AtROPGEF1 from *Arabidopsis thaliana*, OsRacGEF1 from *Oryza sativa*, and PpROPGEF4 from *Physcomitrella patens*. These proteins are known to play critical roles in polar auxin transport during early development, direct interaction with pathogen receptors, and the regulation of apical cell function in moss protonema, respectively (24–26). CrGEF3 and CrGEF4 were placed into Group 1, showing high sequence similarity with *Arabidopsis* AtROPGEF7 and AtROPGEF5, respectively. While AtROPGEF7 is involved in the PLETHORA-dependent maintenance of the root stem cell niche, AtROPGEF5 plays a role in abiotic stress responses in *Arabidopsis* (27, 28). Whereas CrGEF5 was placed into Group 4, along with *Arabidopsis* AtROPGEF14 and *Hordeum vulgare* HvGEF14, which are implicated in cell morphology regulation and ROP activation during pathogen response, respectively (29, 30). On the other hand, phylogenetic analysis of CrGAPs classified them into three distinct groups (Fig. 1 H). CrGAP1 was clustered with PpROPGAP1 and MpROPGAP1, which are known to regulate cell size and coordinate tip growth in *Physcomitrella patens*, and organogenesis in *Marchantia polymorpha*, respectively (31, 32). Whereas CrGAP2 and CrGAP3 were placed in Group 1, showing similarity to *Nicotiana tabacum* NtROPGAP1, and *Arabidopsis* AtROPGAP1 and AtROPGAP4. NtROPGAP1 and AtROPGAP1 have been shown to be involved in pollen tube tip growth (33), whereas AtROPGAP4 plays a role in the plant’s response to oxygen deprivation (34). CrGAP4 was assigned to Group 3, sharing similarity with AtROPGAP3, which is involved in secondary wall pit formation in metaxylem (Oda and Fukuda, 2012). In the case of CrGDIs, phylogenetic analysis categorized them into three groups based on comparison with GDIs from other plants (Fig. 1I). CrGDI1 was uniquely out-grouped into Group 3, while CrGDI2 clustered within Group 1 alongside NtGDI2 from *Nicotiana tabacum* and AtGDI1 from *Arabidopsis thaliana*, both of which are implicated in pollen tube growth (18).

### CrGEF1, CrGAP1, and CrGDI2 Interact with CrROP3 and CrROP5

ROPs function as modulatory switches, cycling between active and inactive states through interactions with regulatory proteins GEFs, GAPs, and GDIs. To elucidate the spatio-temporal regulation of CrROP3 and CrROP5 signalling in modulation of MIA biosynthesis in *C. roseus*, we investigated their potential interactions with RGRPs. We initially conducted an *in-silico* protein-protein interaction analysis using the STRING database, which predicted interactions between CrGEF1, CrGAP1, CrGDI1, and CrGDI2 with both CrROP3 and CrROP5 (SI Appendix, Fig. S5-S7). To experimentally validate the *in silico* interactions, we employed the yeast two-hybrid (Y2H) system. For these assays, the GAL4 activation domain (AD) was fused to the N-termini of each of the 11 RGRP candidates in the pDEST22 vector resulting in prey constructs. To generate bait constructs, GAL4 DNA-binding domain (DBD) was fused to the N-termini of CrROP3 and CrROP5 in the pDEST32 vector. Yeast cells co-transformed with either DBD-CrROP3 or DBD-CrROP5 (bait) along with AD-CrGEF1/2/3/4/5 or AD-CrGAP1/2/3/4, or AD-CrGDI1/2 (prey) successfully grew on double dropout (DDO) selection media (SD/-Trp/-Leu), confirming efficient co-transformation (Fig. 2A-C). Moreover, yeast cells co-transformed with AD-CrGEF1 and DBD-CrROP3 or DBD-CrROP5, AD-CrGAP1 and DBD-CrROP3 or DBD-CrROP5, as well as AD-CrGDI2 and DBD-CrROP3 or DBD-CrROP5, displayed robust growth on quadruple dropout (QDO) selective media (SD/-Trp/-Leu/-His/-Ura), indicating strong protein-protein interactions between these protein-partners (Fig. 2A-C). In contrast, no growth was observed on QDO media for combinations involving AD-CrGEF2/3/4/5 with CrROP3 or CrROP5, AD-CrGAP2/3/4 with CrROP3 or CrROP5, and AD-CrGDI1 with CrROP3 or CrROP5, indicating no interactions between these tested protein-partners (Fig. 2A-C). These results confirm specific interactions of CrGEF1, CrGAP1, and CrGDI2 with CrROP3 or CrROP5. To further validate the interaction results obtained from Y2H assays, we utilized the bimolecular fluorescence complementation (BiFC) system. The candidates, CrGEF1, CrGAP1, and CrGDI2 were cloned in-frame with the split nYFP in the pSPYNE vector, while CrROP3 and CrROP5 were cloned in-frame with the split cYFP in the pSPYCE vector. Tobacco leaves were co-infiltrated with *Agrbacterium* harbouring nYFP-CrGEF1+cYFP-CrROP3, nYFP-CrGEF1+cYFP-CrROP5, nYFP-CrGAP1+cYFP-CrROP3, nYFP-CrGAP1+cYFP-CrROP5, nYFP-CrGDI2+cYFP-CrROP3, and nYFP-CrGDI2+cYFP-CrROP5 constructs. Subsequent confocal microscopy showed clear fluorescence in the cytoplasm and plasma membrane in samples infiltrated with CrGEF1, or CrGAP1, or CrGDI2 with CrROP3 or CrROP5 (Fig. 2D). The observed fluorescence indicated positive interactions of CrGEF1, or CrGAP1, or CrGDI2 with CrROP3 or CrROP5, and corroborated the Y2H results. In contrast, there was no fluorescence observed in leaf samples infiltrated with empty vector control samples co-infiltrated with Agrobacteria harboring pSPYNE-nYFP and pSPYCE-cYFP (Fig. 2D).

**Figure 2.**
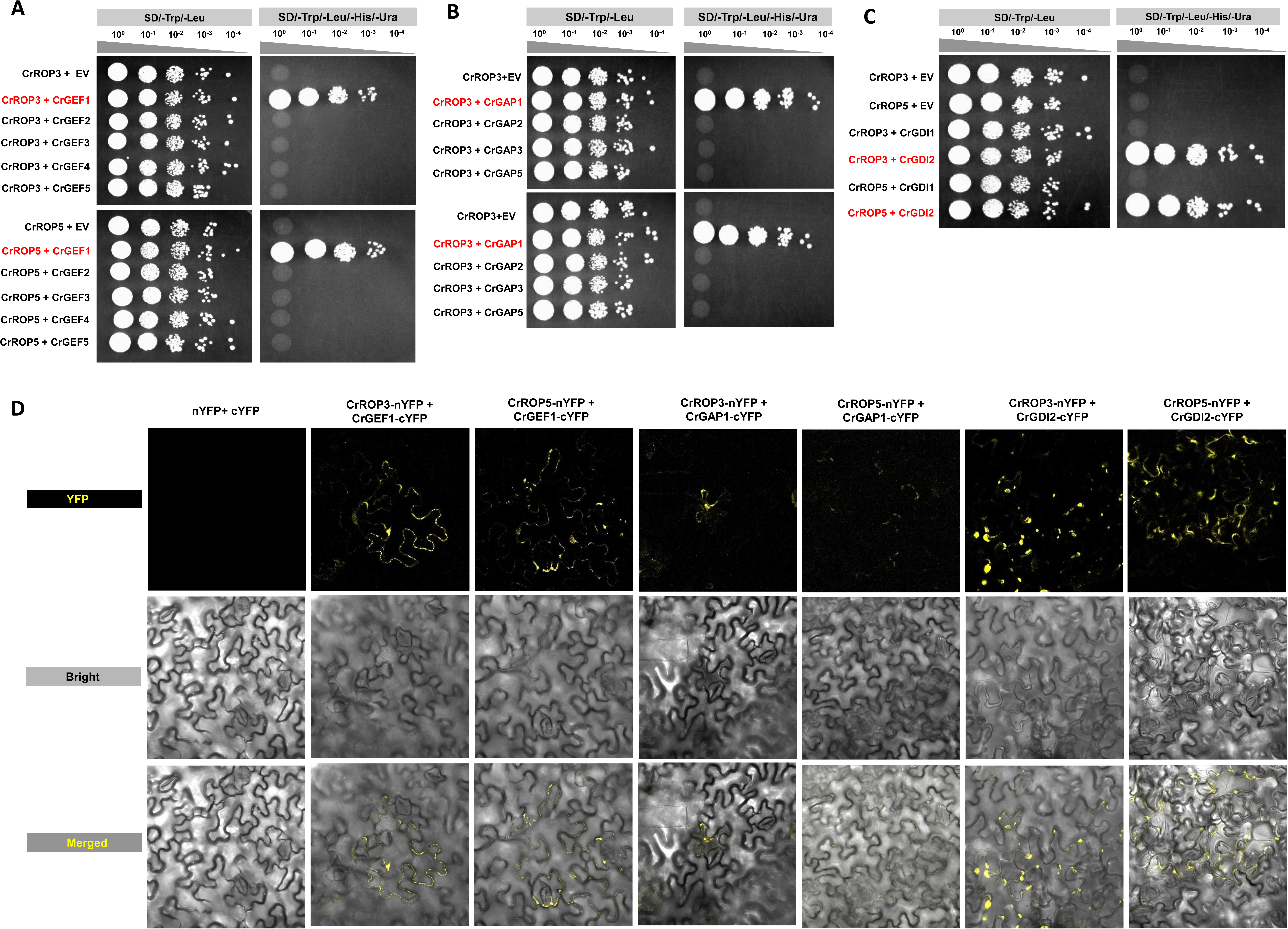
Protein-Protein Interaction Assay of CrGEFs, CrGAPs and CrGDIs with CrROP3 and CrROP5. (A-C) Yeast-two-hybrid assay with pDEST22 (prey) or pDEST32 (bait) constructs containing fusion proteins of the RGRPs and CrROP3 or CrROP5 coupled to respectively, the activator (AD) or binding domain (BD). The empty pDEST22 or empty pDEST32 plasmids were used to check for auto-activation. Transformed yeast grown on the selective -Trp/-Leu (-T -L) medium and interaction verifying -Trp/-Leu/-His/-Ura (-T -L -H) medium. (D) Bimolecular fluorescence complementation (BiFC) assay in tobacco leaf cells testing protein–protein interaction between CrGEF1 and CrROP3, CrGEF1 and CrROP5, CrGAP1 and CrROP3, CrGAP1 and CrROP5, CrGDI2 and CrROP3, CrGDI2 and CrROP5. Reconstitution of yellow fluorescence shows positive interaction in the plasma membrane and cytosol, and absence of fluorescence in the negative control (first panel) shows no interaction between YN and YC alone of YFP.

### Expression Analysis of *CrGEF1*, *CrGAP1* and *CrGDI2*

To investigate the spatio-temporal expression patterns of *CrGEF1*, *CrGAP1*, and *CrGDI2* in various *C. roseus* tissues which include leaves, flowers, siliques, shoots, and roots, transcript levels were quantified using reverse transcriptase-quantitative polymerase chain reaction (qRT-PCR). *CrGEF1*, *CrGAP1* and *CrGDI2* showed expression in all analysed tissues albeit at different levels. *CrGEF1* showed the highest overall expression among the three genes, particularly in aerial tissues. Notably, *CrGEF1* exhibited the greatest expression in young leaves (∼4.5-fold increase), followed by mature leaves (∼3-fold), roots (∼2.5-fold), and exhibited basal expression levels in siliques, shoots, and flowers (Fig 3A). *CrGAP1* was most highly expressed in young leaves (∼1.5-fold), with moderate expression in mature leaves (∼1.2-fold) and siliques (∼1.0-fold), while showing basal expression in roots, shoots, and flowers. Whereas *CrGDI2* was predominantly expressed in young leaves (∼3-fold), followed by mature leaves (∼2.5-fold), roots (∼1.8-fold), and showed minimal expression in siliques, shoots, and flowers (Fig. 3A). The expression patterns observed in the RT-qPCR experiments were consistent with *in silico* expression data derived from the MPGR dataset (SI Appendix, Fig. S8A). Further, the expression of *CrGEF1*, *CrGAP1*, and *CrGDI2* in response to the phytohormone methyl jasmonate (MeJA), a known elicitor of specialized metabolic pathways, was investigated. *C. roseus* geraniol synthase (*CrGES*), a key gene in the MIA pathway known to be induced by MeJA (7), served as a positive control. The results demonstrated that *CrGAP1* was significantly induced by MeJA, peaking at 1 hour with a 5.8-fold increase (Fig. 3B). In contrast, *CrGEF1* and *CrGDI2* showed no significant induction following MeJA treatment at any of the tested time intervals (Fig. 3B). Consistent with these findings, *in silico* analysis using FPKM values from the MPGR database also showed induction of *CrGAP1*, whereas there was no induction of *CrGEF1* and *CrGDI2* in seedlings at different time points (SI Appendix, Fig. S8B).

**Figure 3.**
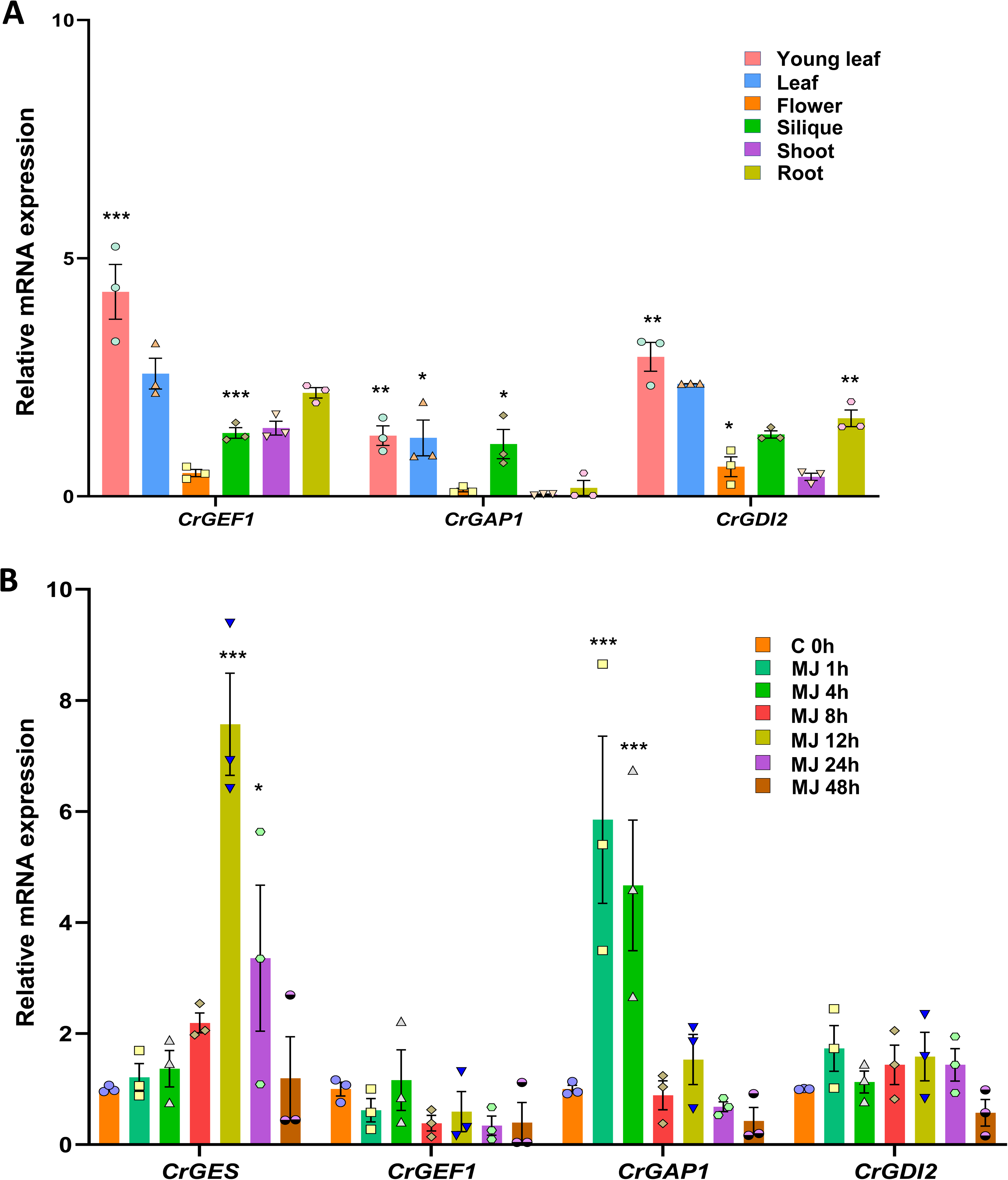
Gene Expression Analysis of *CrGEF1, CrGAP1* and *CrGDI2*. (A) Relative transcript abundance of *CrGEF1, CrGAP1* and *CrGDI2* in different tissues of *C. roseus*. Real time qPCR analysis was performed using total RNA extracted from young leaf, leaf, flower, silique, shoot and root. Expression levels of genes were normalized to the endogenous reference gene *CrN2227*. (B) Expression analysis of *CrGEF1, CrGAP1, CrGDI2* and *CrGES* in response to MeJA. Real time PCR analysis was performed using total RNA extracted from leaves treated with MeJA. Samples were collected at different time intervals. Expression levels of genes were normalized to the endogenous reference gene *CrN2227* and are represented relative to control 0 h, which was set to 1. Statistical significance was calculated with reference to root (A) and 0 h (B). “GraphPad Prism 9” with two-way ANOVA was used to perform statistical analysis:*, *P*=0.033, **, *P*=0.002; ***, *P*<0.001. Each data point represents the mean ± standard error (SE) of three independent experiments with three technical replicates.

### CrGEF1, CrGAP1, and CrGDI2 are Localized in Cytosol/Membrane

RGRPs are pivotal in controlling the activity of ROP proteins, which are primarily active when bound to the plasma membrane in their GTP-bound state and inactive in the cytosol in their GDP-bound form. This cycling between the membrane and cytosol is regulated by GEFs, GAPs, and GDIs. Although RGRPs are typically associated with the plasma membrane, they can also localize to the nucleus and cytosol, influencing ROP signaling across cellular compartments. To investigate the subcellular localization of CrGEF1, CrGAP1, and CrGDI2 in *C. roseus*, we employed a combined approach of *in silico* predictions and experimental validation. *In silico* predictions using tools such as TargetP, WoLF PSORT, ChloroP, and Predotar yielded ambiguous results regarding the localization of CrGEF1, CrGAP1 and CrGDI2 (SI Appendix, Table S2). To obtain more definitive insights, we generated YFP-tagged fusion proteins of CrGEF1, CrGAP1, and CrGDI2 and transiently expressed them in *Nicotiana benthamiana* leaves. To precisely localize CrGEF1, CrGAP1 and CrGDI2, we co-transformed the YFP-tagged fusion proteins with monomeric Red Fluorescent Proteins (mRFP) marker known to be specifically targeted to the plasma membrane and the cytosol. Confocal microscopy analysis of co-transformed leaves revealed a predominant localization of CrGEF1-YFP, CrGAP1-YFP, and CrGDI2-YFP at the cell periphery, indicating their presence at the plasma membrane and within the cytosol (Fig. 4). Furthermore, the clear overlap between the YFP and RFP signals in merged images provided strong evidence for membrane and cytosolic localization of CrGEF1, CrGAP1, and CrGDI2 (Fig. 4). This indicated their crucial role in regulating ROP-GTPase activity at the plasma membrane, while also potentially influencing cytosolic signaling pathways.

**Figure 4.**
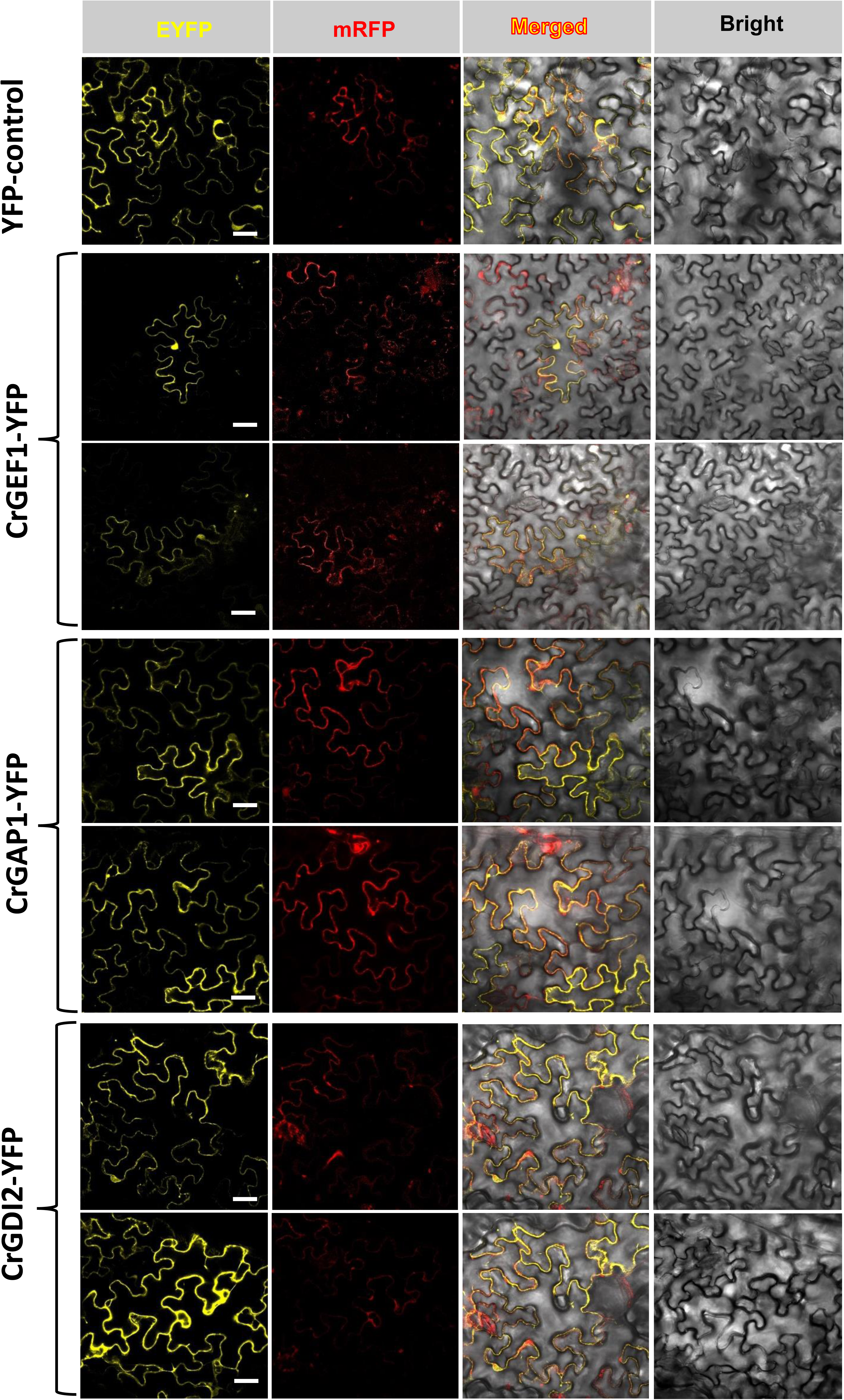
Subcellular Localization of CrGEF1, CrGAP1, and CrGDI2. Confocal laser scanning microscopy of *Nicotiana benthamiana* leaves transformed with CrGEF1-EYFP+mRFP, CrGAP1-EYFP+mRFP, and CrGAP2-EYFP+mRFP co-expression constructs*. N. benthamiana* leaf expressing EYFP+mRFP served as the control. EYFP and mRFP fluorescence are shown in the EYFP and mRFP columns, respectively. The third vertical panel shows the merged image of EYFP and mRFP, indicating colocalization. The fourth and rightmost panel displays the transmission image. Scale bar: 20 µm

### *CrGEF1* Positively Regulates MIA Biosynthesis

To elucidate the *in planta* role of *CrGEF1* in MIAs biosynthesis, we employed a targeted gene silencing approach using virus-induced gene silencing (VIGS). The VIGS constructs were specifically designed to target regions with high sequence dissimilarity from other *CrGEF* isoforms to avoid cross-silencing. The pTRV2-*PDS* construct was used to suppress PDS gene that results in photobleaching, which serves as a visible marker to verify the effectiveness of the VIGS infection and to determine the optimal timing for collecting silenced leaf tissues from *CrGEF1*-vigs plants (SI Appendix, Fig. S9A). qRT-PCR analysis revealed a substantial decrease of *CrGEF1* transcripts (86%) in *CrGEF1*-vigs tissues compared to the EV control (Fig. 5A). However, silencing of *CrGEF1* did not result in any notable visible phenotype (SI Appendix, Fig. S9A). To assess the impact of *CrGEF1* silencing on MIA biosynthesis, we quantified the expression of key genes in the MIA pathway, including TFs (*ORCA3*, *BIS2*, *MYC2*, and *WRKY1*), kinase *MPK3*, and several biosynthetic pathway genes (*DXS*, *GES*, *G10H*, *AS*, *STR*, and *T16H*), which had previously been shown to be affected in *CrROP3* and *CrROP5* silencing background (6). qRT-PCR results revealed a significant reduction in the transcript levels of these TFs, *MPK3*, and pathway genes, with decrease in transcript levels ranging from 40% to 65% in *CrGEF1*-vigs leaf tissues compared to the EV control (Fig. 5B and C). Subsequent metabolite analysis confirmed that this downregulation of key genes related to MIA biosynthesis was associated with a significant decrease of 70% to 80% in the levels of secologanin and downstream MIA such as vindoline, vindolinine, catharanthine, *epi*-vindoline, and ajmalicine (Fig. 5D). To further validate the results of VIGS experiment, *CrGEF1* was overexpressed in *C. roseus* leaves. qRT-PCR analysis showed that overexpression led to a 7-fold increase in *CrGEF1* transcript levels (Fig. 5E), which resulted in an upregulation (ranging from 3- to 6-fold) of the TFs, *MPK3*, and most of the analysed MIA pathway genes, except for *HMGR1* and *AS* (Fig. 5F and G). Further, metabolite profiling revealed a consistent and significant elevation in secologanin and other MIA levels, with an increase of 1.82-fold for secologanin and 1.5- to 1.9-fold for various MIA compared to the EV control (Fig. 5H). Collectively, the results from both VIGS and overexpression experiments demonstrate that *CrGEF1* acts as a positive regulator of MIA biosynthesis by influencing the expression of key TFs and biosynthetic genes, ultimately leading to increased production of MIA.

**Figure 5.**
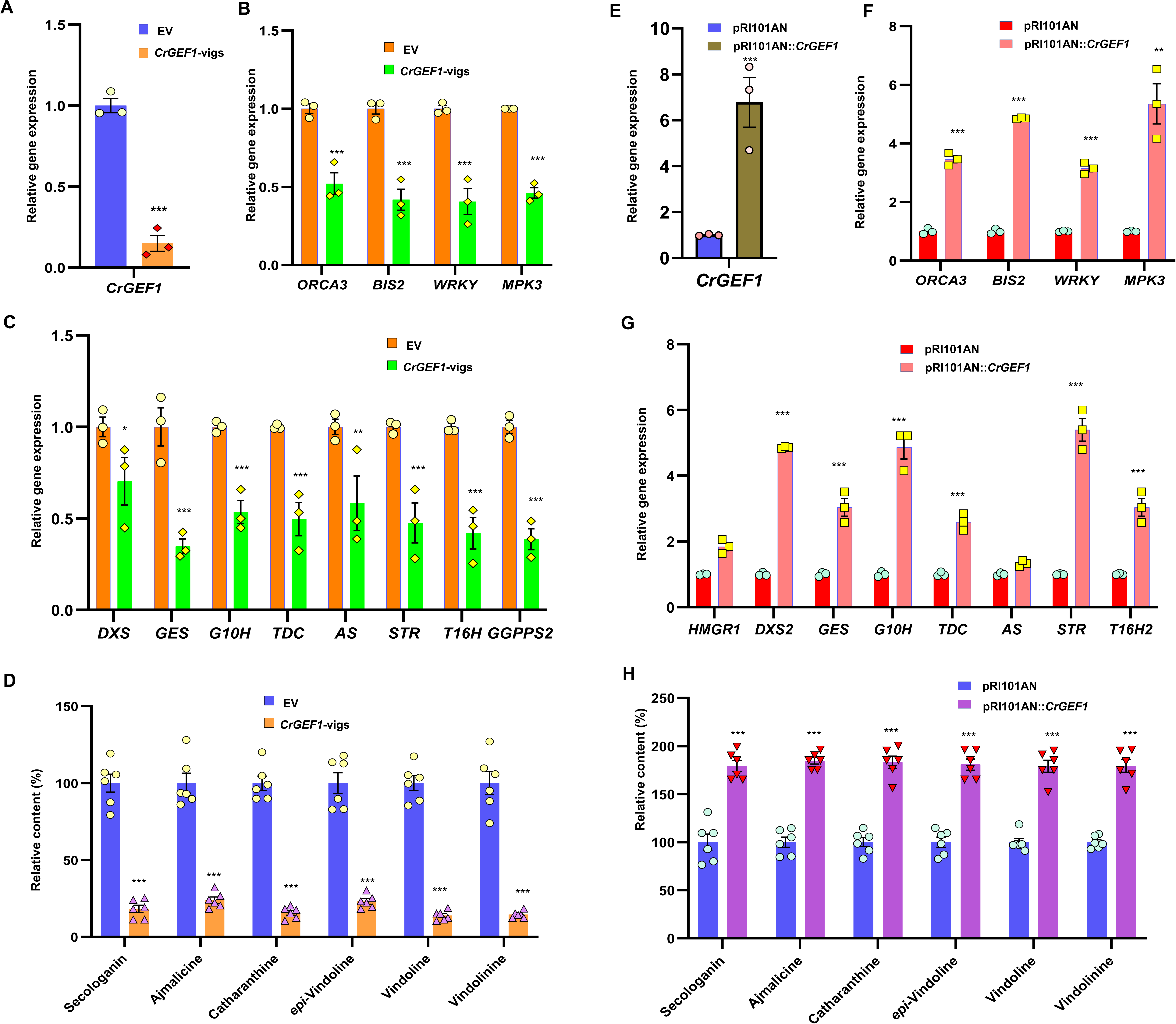
Effect of Silencing and Overexpression of *CrGEF1* on MIA Biosynthesis. Reverse transcriptase-quantitative polymerase chain reaction (RT-qPCR) analysis showing the relative expression of *CrGEF1* in silenced and overexpressed leaves (A & E). Effect of *CrGEF1* silencing and overexpression on the expression of transcription factors (TFs) and pathway genes related to MIA biosynthesis (B, C, F & G) and major MIAs (D & H). Expression levels of genes were normalized to the *CrN2227* endogenous control and were set to 1 in the EV controls to determine the relative reduction in *CrGEF1*-vigs and overexpressed leaves. Levels of different metabolites are represented relative to the EV controls. The results shown are from three (qPCR) and six (metabolites) independent biological replicates with 3 and 2 technical replicates, respectively. Statistics was performed with “GraphPad Prism 9” using one-way ANOVA: ***, *P*<0.001. Error bars indicate mean + SE. Gene abbreviations: *AS*, anthranilate synthase; *BIS2*, basic helix-loop-helix transcription factor 2; *DXS2*, 1-deoxy-D-xylulose 5-phosphate synthase 2; *GES*, geraniol synthase; *G8O/G10H*, geraniol-10-hydroxylase/ 8-oxidase;; *MPK3*, MAP kinase 3; *ORCA3*, octadecanoid responsive *Catharanthus* AP-2 domain 3; *STR*, strictosidine synthase; *T16H2*, tabersonine 16-hydroxylase 2; *TDC*, tryptophan decarboxylase; *WRKY*, WRKY transcription factor.

### *CrGAP1* and *CrGDI2* Negatively Regulate MIA Biosynthesis

To investigate the roles of *CrGAP1* and *CrGDI2* in modulation of MIA biosynthesis, we downregulated their expression in *C. roseus* by VIGS. VIGS constructs targeting unique regions of *CrGAP1* and *CrGDI2* were carefully designed to avoid unintended silencing of related isoforms. qRT-PCR analysis revealed a substantial knockdown of *CrGAP1* and *CrGDI2* transcripts, with reductions of 77% and 79%, respectively (Fig 6A and I), in the newly emerging leaves of *CrGAP1*-vigs and *CrGDI2*-vigs samples relative to the EV control. Notably, this reduction in *CrGAP1* expression led to a pronounced upregulation of multiple transcription factors, MPK3, and most MIA biosynthetic genes, with increases ranging between 4 to 10-fold (except for *TDC, AS, T16H, and GGPS2*, which did not show significant enhancement) (Fig. 6B and C). Similarly, *CrGDI2*-vigs samples exhibited a corresponding increase in gene expression, also ranging from 4 to 10-fold, with the same exceptions (*TDC*, *AS, STR, T16H*, and *GGPS2* which did not show significant enhancement) (Fig. 6 J and K). This enhanced gene expression was accompanied by a significant increase in the accumulation of secologanin and other MIA. Specifically, silencing of *CrGAP1* and *CrGDI2* resulted in a 1.6-to-1.82-fold increase in secologanin levels and a 2-to-3.7-fold elevation in the levels of downstream MIA, including vindoline, catharanthine, *epi*-vindoline, vindolinine, and ajmalicine (Fig. 6D and L). To further validate these findings, *CrGAP1* and *CrGDI2* were overexpressed in *C. roseus* leaves, which led to increase in the transcript levels of *CrGAP1* (8.6-fold) and *CrGDI2* (up to 9.8 fold) (Fig. 6E and M). Overexpression of *CrGAP1* and *CrGDI2* resulted in a significant downregulation of all analysed TFs, *MPK3*, and MIA pathway genes, with reductions ranging from 40% to 70% (except for *TDC, AS, STR, T16H*, which did not show significant reduction) (Fig. 6F,G,M and O). Subsequent metabolite analysis confirmed that this transcript downregulation was associated with a marked decrease in the levels of secologanin and other MIA, ranging from 60 to 80% compared to EV controls (Fig. 6H and P). These results showed that, in stark contrast to *CrGEF1*, *CrGAP1* and *CrGDI2* act as negative regulators of MIA biosynthesis, complementing the positive regulatory role of *CrGEF1* thus highlighting the complex regulatory network governing MIA production in *C. roseus*.

**Figure 6.**
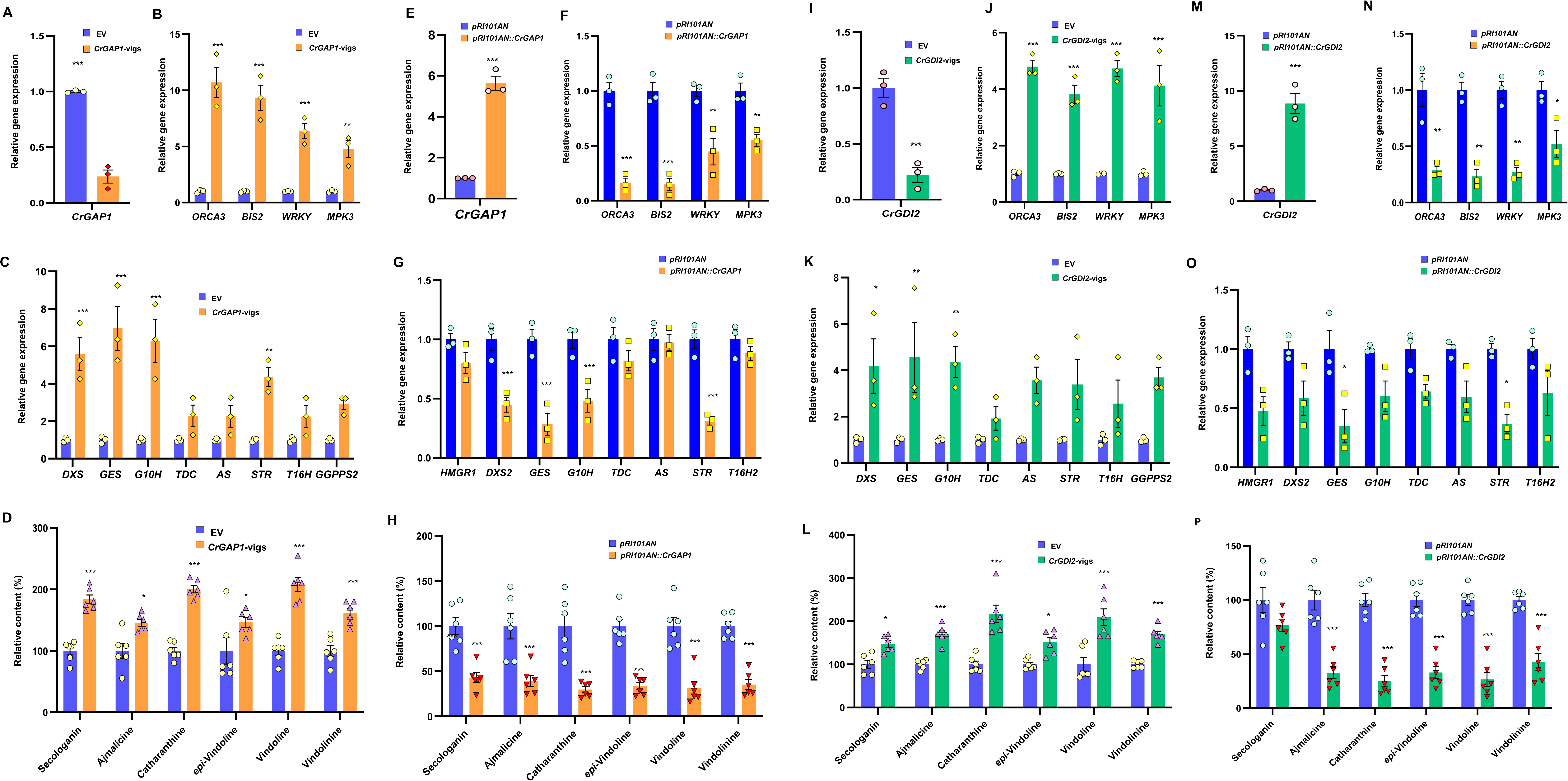
Effect of Silencing and Overexpression of *CrGAP1* and *CrGDI2* on MIA Biosynthesis. Reverse transcriptase-quantitative polymerase chain reaction (RT-qPCR) analysis showing the relative expression of *CrGAP1* and CrGDI2 in silenced and overexpressed leaves (A,E,I & M). Effect of *CrGAP1* and *CrGDI2* silencing and overexpression on the expression of transcription factors (TFs) and pathway genes related to MIA biosynthesis (B, C, F, G, J, K, N, & O) and major MIAs (D, H, L, & P). Expression levels of genes were normalized to the *CrN2227* endogenous control and were set to 1 in the EV controls to determine the relative reduction in *CrGAP1*-vigs and *CrGDI2*-vigs and overexpressed leaves. Levels of different metabolites are represented relative to the EV controls. The results shown are from three (qPCR) and six (metabolites) independent biological replicates with 3 and 2 technical replicates, respectively. Statistics was performed with “GraphPad Prism 9” using one-way ANOVA: ***, *P*<0.001. Error bars indicate mean + SE. Gene abbreviations: *AS*, anthranilate synthase; *BIS2*, basic helix-loop-helix transcription factor 2; *DXS2*, 1-deoxy-D-xylulose 5-phosphate synthase 2; *GES*, geraniol synthase; *G8O/G10H*, geraniol-10-hydroxylase/ 8-oxidase; *MPK3*, MAP kinase 3; *ORCA3*, octadecanoid responsive *Catharanthus* AP-2 domain 3; *STR*, strictosidine synthase; *T16H2*, tabersonine 16-hydroxylase 2; *TDC*, tryptophan decarboxylase; *WRKY*, WRKY transcription factor.

### N-terminal Domain of CrGEF1, and C-terminal Domain of CrGAP1 and CrGDI2 are Needed for Interaction with CrROP3 and CrROP5

Silencing and overexpression experiments collectively demonstrated the regulatory role of CrGEF1, CrGAP1, and CrGDI2 in modulating MIA biosynthesis in *C. roseus* leaves. The presence of variable N- and C-terminal regions in CrGEF1, CrGAP1, and CrGDI2, which often contain crucial regulatory motifs or signal sequences, prompted us to investigate whether these specific domains have any regulatory role in interactions of CrGEF1, CrGAP1 and CrGDI2 with CrROP3 or CrROP5 proteins, and thus on MIA biosynthesis. To investigate this, we generated a series of truncated versions of CrGEF1, CrGAP1, and CrGDI2, each lacking either the variable N-terminal or C-terminal regions and also generated mutants retaining only conserved domains such as the PRONE domain in CrGEF1, the CRIB and GAP domains in CrGAP1, and the Rho-GDI domain in CrGDI2 (Fig. 7A-C). These truncated and mutant versions were cloned into pDEST22 vector (prey) for Y2H assays to check their interactions with CrROP3 and CrROP5 proteins which were cloned in pDEST32 vector (Bait). Y2H analyses for CrGEF1 truncated version, showed that the C-terminal deletion construct (ΔCCrGEF1) co-transformed with CrROP3 or CrROP5 showed growth in DDO (SD/-Leu/-Trp) and QDO selection media (SD/-Leu/-Trp/-His/-Ura) indicating its interaction with either CrROP3 or CrROP5. Conversely, the N-terminal deletion construct (ΔNCrGEF1) and the construct containing only the PRONE domain (PRONECrGEF1) when co-transformed with CrROP3 or CrROP5 showed growth in DDO selection media but failed to grow in QDO selection media (Fig. 7A), suggesting that the N-terminal region contains crucial elements necessary for stabilizing the interaction or enabling the PRONE domain to adopt a conformation conducive to binding. In the case of CrGAP1, five distinct truncations were generated to systematically evaluate the contributions of its domains to ROP interaction. Only the N-terminal deletion construct (ΔNCrGAP1) retained the ability to interact with CrROP3 and CrROP5, implying that the N-terminal region may contain inhibitory elements that, when removed, enhance the interaction potential of the GAP domain (Fig. 7B). Whereas other truncated constructs containing only the CRIB and GAP domains, C-terminal deletion, CRIB and GAP domain combined were unable to interact with the ROPs (Fig. 7B). Similarly, for CrGDI2, the N-terminal deletion construct (ΔNCrGDI2) was able to interact with both CrROP3 and CrROP5, while the C-terminal deletion (ΔCCrGDI2) and the construct harbouring only the GD1 domain (GDICrGDI2) failed to show any interaction (Fig. 7C). Together, these results indicate that the N-terminal domain of CrGEF1, and the C-terminal domains of CrGAP1 and CrGDI2 are essential for their interactions with CrROP3 and CrROP5.

**Figure 7.**
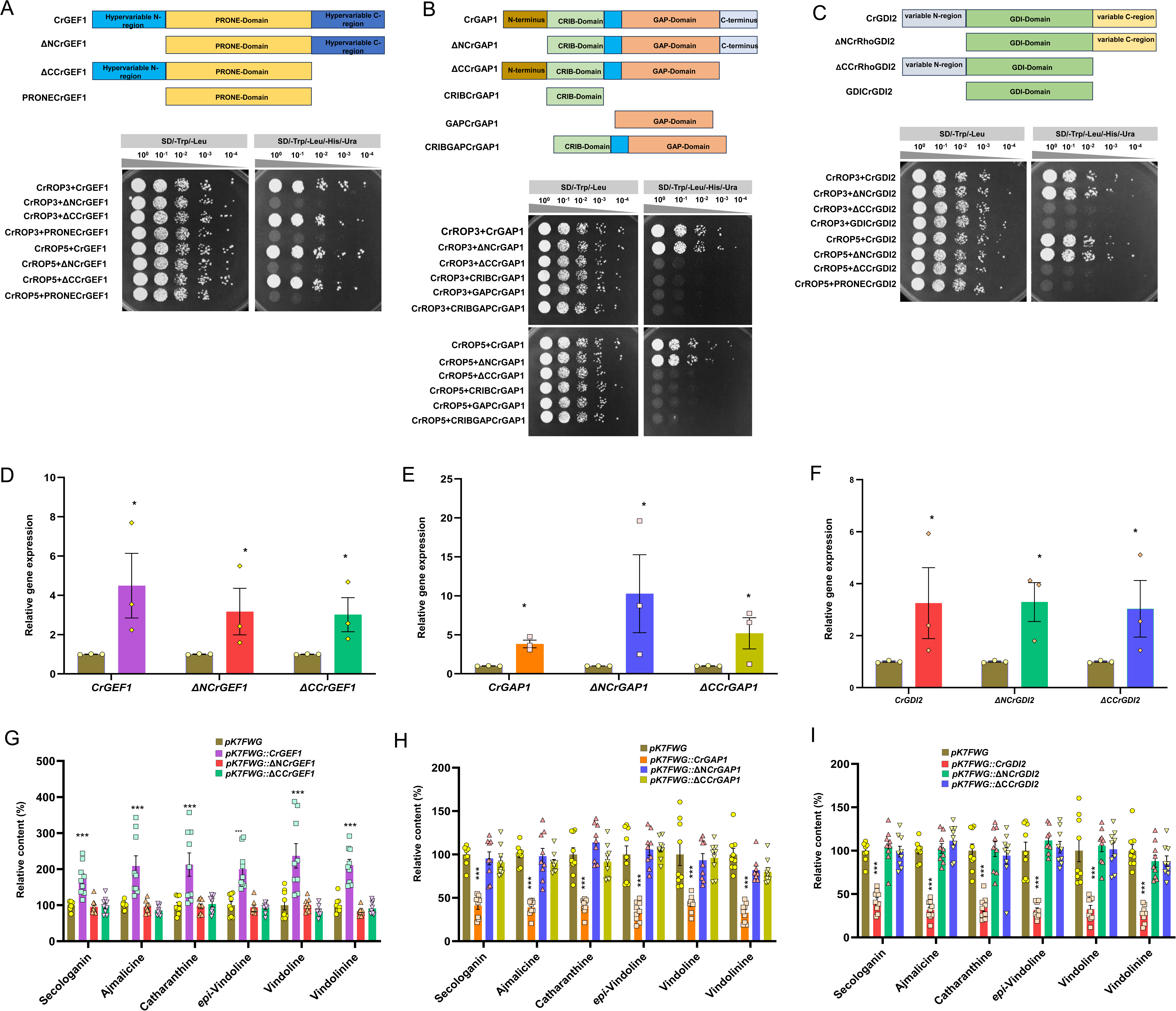
Interaction of Mutant RGRPs with CrROP3 or CrROP5 and its Effect on MIA Biosynthesis. (A, B, and C) Upper panel: Schematic representation of different truncated versions of CrGEF1, CrGAP1, and CrGDI2. Lower panel: Co-transformation screening in yeast two-hybrid (Y2H) assays showing interactions of truncated versions of CrGEF1, CrGAP1, and CrGDI2 with CrROP3 and CrROP5 (D, E and F) Relative gene expression of *CrGEF1*, *ΔNCrGEF1*, *ΔCCrGEF1*, *CrGAP1*, *ΔNCrGAP1, ΔCCrGAP1, CrGDI2*, *ΔNCrGDI2* and *ΔCCrGDI2* in *C. roseus* leaves agroinfiltrated with wild-type and mutant RGRP constructs, respectively, in comparison to EV control. Expression levels of genes were normalized to the endogenous reference gene CrN2227 and are represented relative to the pK7FWG control, which was set to 1. (G, H and I) Effect of overexpression of mutant RGRPs on accumulation of MIA. Levels of MIAs were analyzed by HPLC and are expressed in % relative to pK7FWG control. Statistics was performed with “GraphPad Prism 9” using two-way ANOVA: ***, P<0.001. Error bars indicate mean + SE.

### Overexpression of Truncated RGRP Interacting Partners of CrROP3/CrROP5 do not Affect MIA Biosynthesis

As the truncated ΔCCrGEF1, ΔNCrGAP1, and ΔNCrGDI2 demonstrated interactions with CrROP3 and CrROP5, we sought to determine whether these truncated forms are sufficient to retain the functional roles of their wild-type counterparts in modulation of MIA biosynthesis. To address this, agroinfiltration experiments were conducted in *C. roseus* leaves using constructs for both the wild-type and their corresponding truncated mutants. Specifically, leaves were agroinfiltrated with constructs for wild-type CrGEF1, CrGAP1, and CrGDI2, as well as mutants ΔCCrGEF1, ΔNCrGAP1 and ΔNCrGDI2. These were compared against the vector control (EV) and negative controls (ΔNCrGEF1, ΔCCrGAP1, and ΔCCrGDI2), which lacked regions essential for interaction with CrROP3 and CrROP5. Transcript analysis using quantitative PCR revealed that transcripts of both wild-type and mutant RGRPs were successfully expressed, with their levels elevated by 6.56-fold to 8.6-fold compared to the EV control (Fig. 7D-F). As observed previously, overexpression of wild-type *CrGEF1* significantly increased MIA levels (1.5 to 2.17-fold compared to EV), while overexpression of *CrGAP1* and *CrGDI2* reduced MIA content (Fig. 7G,H and I). In contrast, the truncated mutants, *ΔCCrGEF1*, *ΔNCrGAP1*, and *ΔNCrGDI2*, failed to induce any significant changes in MIA accumulation, indicating that the deletion of critical N- or C-terminal domains disrupted their ability to modulate MIA biosynthesis. Additionally, overexpression of RGRP mutants *ΔNCrGEF1*, *ΔCCrGAP1*, and *ΔCCrGDI2* (controls) also had no effect on MIA levels (Fig. 7D-F). These results provide compelling evidence that the specific domains identified as critical for interaction of CrGEF1 or CrGAP1 or CrGDI2 with CrROP3 or CrROP5 in Y2H analysis, are not solely sufficient for their biological function *in planta*. This indicated that both N- and C-terminal domains together are involved in enabling these proteins to exert their full regulatory role in MIA biosynthesis.

## Discussion

Results reported in this study have uncovered an intricate signalling network underlying MIA biosynthesis, involving activation and inactivation of specific ROP-GTPases CrROP3 and CrROP5, by distinct RGRPs CrGEF1, CrGAP1 and CrGDI2 (Fig. 8). Our results demonstrate that the CrGEF1-CrROP3/CrROP5 module positively influences the expression of TFs and pathway genes, enhancing MIA biosynthesis, whereas the CrGAP1-CrROP3/CrROP5 and CrGDI2-CrROP3/CrROP5 modules negatively affect the expression of these genes, resulting in reduced MIA biosynthesis (Fig. 5 and Fig. 6). Similar to ROP-GTPases which are encoded by multigene family(6), *C. roseus* contains expanded repertoire of RGRPs, including five ROP-GEFs, four ROP-GAPs, and two ROP-GDIs distributed across four chromosomes (Supplementary Fig. 1). This expansion likely reflects an evolutionary diversification for controlling complex signalling processes in response to different environmental and developmental cues, as has been observed from non-vascular to vascular plants, highlighting a common evolutionary trajectory across plant species(11, 12, 15, 31). Our phylogenetic analysis showed that *C. roseus* RGRPs cluster with homologous proteins of other plants having varied functions, suggesting that the expansion and diversification of RGRPs have been critical for the evolution of specialized functions in plant signalling networks (Fig. 1G-I).

**Figure 8.**
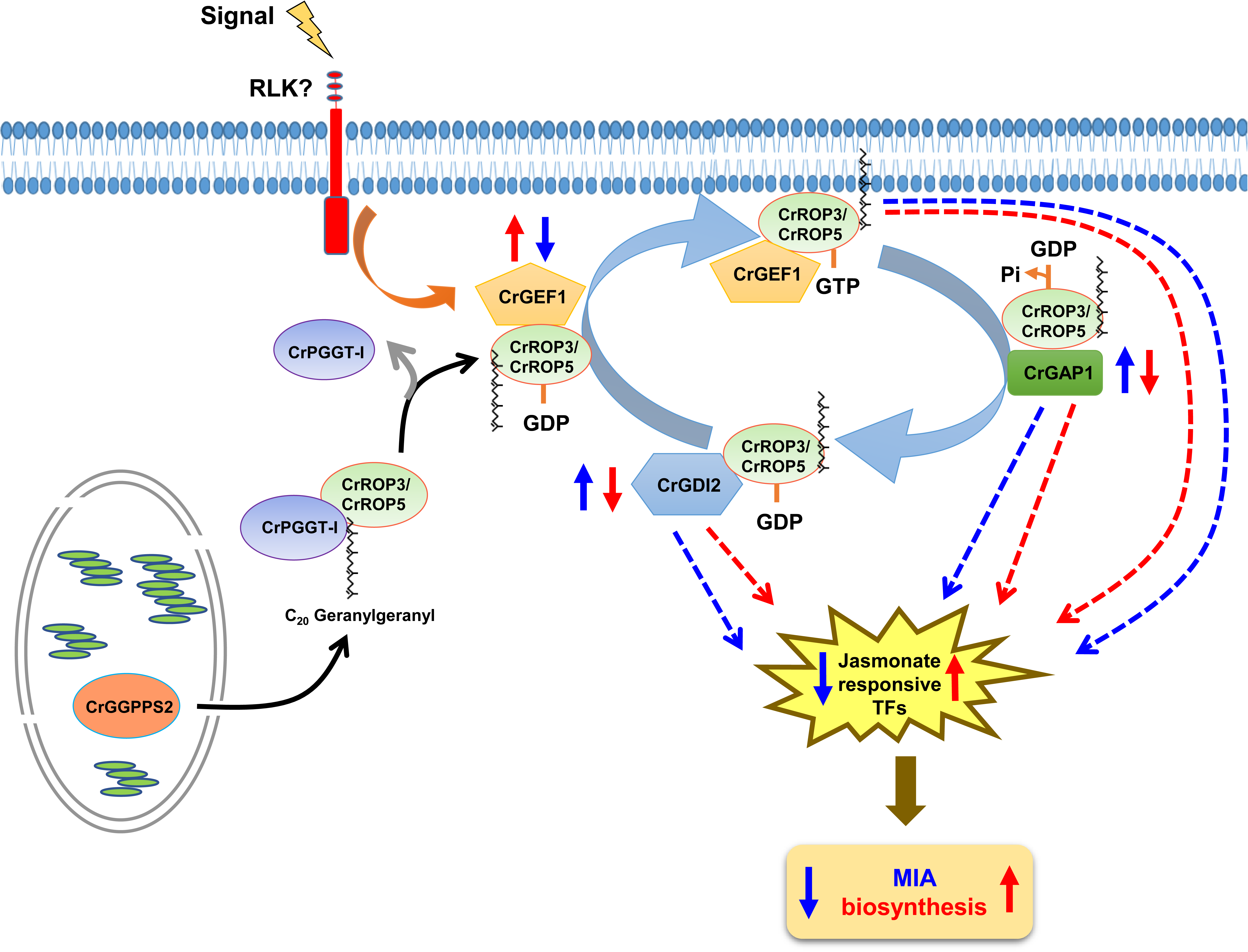
Proposed Model of ROP/RGRP-mediated Modulation of MIA Biosynthesis in C. roseus. CrPGGT-I interacts with CrROP3 or CrROP5, and geranylgeranylates them by utilizing GGPP made by plastidial CrGGPPS2. Further, CrROP3 and CrROP5 interact with RGRPs CrGEF1, and CrGAP1 and CrGDI2 to bring about positive and negative modulation of MIA biosynthesis, respectively. Down-regulation of *CrGEF1* reduces the activation of CrROP3/5 from GDP-bound inactive form into their active GTP-bound forms thereby negatively affecting MIA biosynthesis. Similarly, downregulation of *CrGAP1* reduce the GTPase activity of CrROP3/5 thereby positively affecting MIA biosynthesis. *CrGDI2* silencing reduces the sequestration of CrROP3/5 from membrane to cytosol thereby positively affecting MIA biosynthesis.

ROP-GTPases CrROP3 and CrROP5 belonging to type-II ROPs, are involved in the modulation of MIA biosynthesis(6). They are functionally divergent from type-I ROPs, which are generally involved in basic cellular functions such as cell growth and development (12, 35). The involvement of CrROP3 and CrROP5 in MIA biosynthesis suggests that type-II ROPs have evolved unique roles to support secondary metabolism. Though both ROPs and RGRPs in plants are of multigene family, preferential interactions between distinct ROP and RGRP candidates has been shown to selectively affect defined processes like cell polarity, morphogenesis, hormonal regulation and immune response providing functional specificity to ROP signalling (12, 16, 20). Accordingly, selective interactions of RGRPs CrGEF1, CrGAP1, and CrGDI2, with CrROP3 and CrROP5, that have been shown to affect MIA formation, indicated their role in modulation of signalling in a spatio-temporal manner to control MIA biosynthesis (Fig. 2)(6). Indeed, *in planta* VIGS and overexpression of genes encoding CrGEF1, CrGAP1, and CrGDI2, that showed interaction with CrROP3 and CrROP5, corroborated their role in controlling MIA biosynthesis. Silencing *CrGEF1* led to a significant reduction in the expression of TFs and pathway genes related to MIA biosynthesis, resulting in decreased MIA content. Conversely, overexpression of *CrGEF1* led to enhanced MIA biosynthesis, underscoring its role as an essential activator of ROP signalling in the context of MIA production (Fig. 5). Silencing of *CrGAP1* and *CrGDI2*, on the other hand, led to upregulation of TFs and pathway genes resulting in enhanced MIA levels, while their overexpression had the opposite effect (Fig. 6). This demonstrated that CrGAP1 and CrGDI2 act as negative regulators, attenuating ROP signalling to avoid excessive metabolite accumulation. Such regulatory flexibility has been shown in *Arabidopsis*, where distinct ROP-GEFs, ROP-GAPs and ROP-GDIs modulate ROP signalling in a context-dependent manner, controlling signalling intensity based on tissue type and environmental conditions (17, 18, 36, 37). Moreover, *CrGEF1, CrGAP1* and *CrGDI2* exhibited co-expression with *CrROP3* and *CrROP5*, and *PGGT-I* supporting their co-ordinated role in MIA biosynthesis (Fig. 3) (6, 7).

The fact that CrGEF1, CrGAP1 and CrGDI2 were localized to plasma membrane and cytosol, and they interact with CrROP3 and CrROP5, showed that these specific RGRPs regulate their function both at PM and cytosol (Fig. 2,4 and SI Appendix, Fig. S13), where ROP-GTPases are known to execute their signaling by relaying signals to downstream effectors such as ROP-Interactive CRIB Motif-containing proteins (RICs) and Rho GTPase-Interacting Proteins (RIPs) which further contribute to the specificity of ROP signaling by translating ROP activation into cellular responses (Fig. 2 and Fig. 4) (12, 16). While the plasma membrane localization likely enables the precise modulation of ROP activity at membrane sites, crucial for efficient signal transduction in response to cellular cues, the cytosolic localization has role in sequestering inactive ROPs, allowing for controlled release and activation of ROPs upon appropriate signaling. Such spatial control mechanisms have been reported in *Arabidopsis*, where ROP-GEFs localize to specific cellular regions to modulate growth (16, 37). Further complexity in RGRPs regulation has also been reported, where it was shown that receptor-like kinases (RLKs) and cytosolic kinases, such as MAP kinases, calcium-dependent protein kinases (CDPKs), and BRASSINOSTEROID-INSENSITIVE2 (BIN2), can influence the activity of RGRPs through phosphorylation, introducing an additional layer of regulation (38, 39). Though mutant ΔNCrGEF1, ΔCCrGAP1, and ΔCCrGDI2 lacking N- or C-terminal motifs, respectively, were able to efficiently interact with CrROP3 and CrROP5, they failed to modulate the MIA biosynthesis when overexpressed unlike their wild type counterparts (Fig. 7). This indicated the importance of specific conserved non-catalytic terminal domains in CrRGRPs are essential for localization and interaction specificity in ROP signaling (15, 17, 23, 40).

Overall, this comprehensive characterization of CrRGRPs uncovers a feedback regulatory mechanism involving CrGEF1, CrGAP1, and CrGDI2 in modulating MIA biosynthesis through their preferential interactions with CrROP3 and CrROP5 in *Catharanthus roseus*. The specialization of these regulatory proteins and their selective interactions with distinct ROPs underline a finely tuned signalling network that regulates MIA production. Our findings also highlight the evolutionary significance of RGRP expansion and functional specialization, suggesting that these proteins have developed specific roles within ROP signalling pathways to support secondary metabolism. Further, this study underscores the potential of leveraging these regulatory mechanisms in metabolic engineering strategies aimed at enhancing the production of valuable plant metabolites. Furthermore, this study lays a foundation for future investigations into the ROP-RGRP module and other effectors or signalling components of secondary metabolism, particularly their roles in upstream signal perception and downstream signal relaying. By elucidating the molecular basis of these ROP signalling networks, one can better understand the regulation of jasmonate signalling a key pathway in MIA production in *C. roseus* and secondary metabolism in plants.

## MATERIALS AND METHODS

### Plant material, tissue collection and MeJA treatment

*Catharanthus roseus* cv. Dhawal (National Gene Bank, CSIR-CIMAP, Lucknow, India) plants were grown in a controlled environment at a temperature of 22-25°C with a 16-hour light and 8-hour dark cycle. MeJA treatment was performed as described in our previous study (6). Briefly, the excised first pair of leaves were dipped in 200 µM MeJA solution or water (control) containing the same amount of DMSO without MeJA. Samples were collected at different time points and stored at −80 °C for further use. For tissue-specific expression analysis, roots, stems, leaves, siliques, and flowers were collected separately from 8-wk-old plants, flash frozen in liquid nitrogen and stored at −80 °C until further analysis. For VIGS and transient overexpression experiments, seeds were germinated and grown as mentioned above. In the VIGS experiment, fully developed first and second pairs of leaves from six-leaf-staged plants were utilized. In transient overexpression experiment, fully developed first- and second-pair leaves from 6- to 8-week-old plants were used for gene expression and metabolite analyses.

### Mining of CrRGRPs from the MPGR database

To identify potential genes encoding CrRGRPs in *C. roseus*, a search was carried out in the MPGR (http://medicinalplantgenomics.msu.edu) database initially by using RGRPs of other plants as query and then later by blast search against *C. roseus* genome sequence (21). Retrieved sequences were then stringently filtered. Sequences with low or no FPKM value and short transcript lengths were excluded. This resulted in a final set of 11 CrRGRP candidates: 5 encoding GEFs, 4 encoding GAPs, and 2 encoding GDIs.

### Phylogenetic analysis

For phylogenetic analysis, amino acid sequences of characterized RhoGEFs, RhoGAPs and RhoGDIs from different plant species were retrieved from the National Center for Biotechnology Information (NCBI) data base (www.ncbi.nlm.nih.gov). The deduced amino acid sequences of CrGEFs, CrGAPs and CrGDIs were compared with other RhoGEFs, RhoGAPs and RhoGDIs proteins obtained from the NCBI database. The phylogenetic tree was constructed by neighbour-joining method using MEGA11 software (41). Multiple sequence alignment of CrGEFs, CrGAPs and CrGDIs proteins was performed using ClustalW (42) with default parameters through https://www.genome.jp/tools-bin/clustalw.

### Yeast two-hybrid assay

For yeast two hybrid assay, coding sequences of *CrGEF1-5*, *CrGAP1-4* and *CrGDI1-2* were PCR amplified from leaf cDNAs using Phusion high-fidelity DNA polymerase (Thermo Fisher Scientific) and gene-specific oligonucleotides, and the amplicons were cloned in-frame with the activation domain of pDEST22 (prey) vector to generate *CrGEF1-5-*AD, *CrGAP1-4* -AD *CrGDI1-2 -*AD, truncated CrGEF1-AD, truncated CrGAP1-AD, and truncated CrGDI2-AD constructs. Similarly, coding sequences of *CrROP3* and *CrROP5* were amplified and cloned in-frame with the DNA binding domain of pDEST32 (bait) vector to generate *CrROP3-*DBD and *CrROP5-*DBD constructs. The resulting constructs were co-transformed into MaV203 yeast strain and positive colonies screen through colony PCR were selected and cultured on double dropout (DDO) supplement (SD-Leu/-Trp) for 2–3 days, then co-transformants were shifted on to quadruple QDO supplement (SD-Leu/-Trp/-His/-Ura) to test for possible interactions.

### Bimolecular fluorescence complementation assay

The ORF of *CrGEF1*, *CrGAP1* and *CrGDI2* were cloned into pSPYCE-nYFP to generate the nYFP-CrGEF1, nYFP-CrGAP1 and nYFP-CrGDI2 vectors, respectively. Likewise, the CDS of *CrROP3* and *CrROP5* were cloned into pSPYNE-*cYFP* to generate the cYFP-CrROP3 and cYFP-CrROP5 vectors, respectively. The resulting constructs were transferred into *A. tumefaciens* strain GV3101 using freeze thaw method. Positive colonies were selected and cultured at 28°C for 24 h, then the bacteria were centrifuged and resuspended in infiltration medium (10 mM MES, pH 5.6, 10 mM MgCl_2_, and 200 μM acetosyringone) to a final OD600 of 0.6 – 0.8 and the agroinfiltration was performed on leaves of 5- to 6-week-old *N. benthamiana* plant using needleless syringe. *A. tumefaciens* with p19 (suppressor of RNA silencing) construct was mixed with Agrobacterium harboring split-nYFP constructs or split-cYFP in a 1:1:1 ratio prior to infiltration. The agroinfiltrated plants were kept in dark for 48 hours then kept in a growth chamber for at least 12-24 h and fluorescent signals were observed under a Carl Zeiss LSM880 laser scanning confocal microscope using a 63× oil immersion objective with a numerical aperture of 1.4. YFP fluorescence was excited at 514 nm and detected in the range of 525 to 562 nm.

### Analysis of subcellular localization

The subcellular localization of CrGEF1, CrGAP1 and CrGDI2 was determined by generating YFP fusion protein using the pGWB441 vector. *CrGEF1, CrGAP1* and *CrGDI2* were cloned into pGWB441 to express them as C-terminal EYFP-tagged proteins. The resulting pGWB441::*CrGEF1-YFP*, pGWB441::*CrGAP1-YFP* and pGWB441::*CrGDI2-YFP* constructs were mobilized into *Agrobacterium tumefaciens* (GV3101) using the freeze-thaw method. For transient expression of the fusion proteins in *N. benthamiana* leaves, *Agrobacterium* suspension was prepared in infiltration buffer (10 mM MES, pH 5.6, 10 mM MgCl2, and 200 μM acetosyringone), and the agroinfiltration was performed on leaves of 5- to 6-week-old plants. *A. tumefaciens* with p19 (suppressor of RNA silencing) construct was mixed with *Agrobacterium* harboring overexpression fusion-YFP constructs or EV and mRFP in a 1:1:1 ratio prior to infiltration. After 48 to 56 h of agroinfiltration, leaf discs were analyzed, and images were captured under a Carl Zeiss LSM880 laser scanning confocal microscope using a 63× oil immersion objective with a numerical aperture of 1.4. YFP fluorescence was excited at 514 nm and detected in the range of 525 to 562 nm, whereas RFP was excited at 561 nm and detected in a range of 570 to 652 nm.

### Generation of silencing and overexpression constructs

The pTRV1 and pTRV2 vectors used for generating VIGS constructs were procured from The Arabidopsis Information Resource (TAIR), USA. The 450-500 bp fragments of *CrGEF1, CrGAP1,* and *CrGDI2* were amplified from leaf cDNA by PCR using gene specific primers having *Eco*RI restriction site at 5′ end of each primer (SI Appendix, Table S3). The resulting amplicon was purified and sub-cloned into pJET1.2/vector and sequences were confirmed by nucleotide sequencing. The confirmed fragment was then cloned into pTRV2 vector digested with *EcoR*I. The resulting pTRV2::*CrGEF1* or *CrGAP1* or *CrGDI2* plasmid was confirmed by digestion. For generation of overexpression constructs, the ORF of *CrGEF1, CrGAP1,* and *CrGDI2* were PCR amplified using leaf cDNA with specific forward and reverse primers (SI Appendix, Table S3). A truncated CrGEF1, CrGAP1 and CrGDI2 constructs lacking terminal region was generated by deleting N and C terminal variable region. The amplicons were cloned into pJET1.2/ cloning vector for sequence confirmation, and then sub-cloned into *BamHI* and *EcORI* sites of pRI101AN binary vector under the control of the 35S promoter of Cauliflower mosaic virus (CaMV) to form pRI101AN::*CrGEF1* or pRI101AN::*CrGAP1* pRI101AN::*CrGDI2* constructs (SI Appendix, Fig. S12C). The VIGS and overexpression constructs were then mobilized into *Agrobacterium tumefaciens* GV3101 competent cells by freeze thaw method.

### VIGS and Transient Overexpression

VIGS and transient overexpression was performed according to the Kumar *et. al.*, 2020. For silencing, briefly the overnight grown *Agrobacterium* cultures harboring pTRV1 and pTRV2::*CrGEF1* or pTRV2::*CrGAP1* or pTRV2::*CrGDI2* or pTRV2::*CrPDS* or pTRV2 (EV) constructs were resuspended in MES buffer (10 mM MES, pH 5.6, 10 mM MgCl2, and 200 μM acetosyringone) to a final OD_600_ of 1.6 and incubated at 28 °C for 3 h in shaker incubator and mixed in 1:1 ratio before infiltration. Plants were infected by pricking below the apical meristem using a dissecting needle and kept in dark for 48 h. Plants were shifted to a growth chamber (22°C, 75% humidity, 16-8h light cycle). Leaves from pTRV2::*CrGEF1,* pTRV2::*CrGAP1,* pTRV2::*CrGDI2,* and pTRV2 infected plants were harvested 3-4 weeks post infection, when albino phenotype developed in the first 2 pairs of leaves in pTRV2::*CrPDS* infected plants, and stored in -80°C for further use. Transient overexpression experiments were performed using *Agrobacterium tumefaciens* harboring the constructs pRI101AN::*CrGEF1*, pRI101AN::*CrGAP1*, pRI101AN::*CrGDI2*, or pRI101AN (empty vector control). For truncated protein versions, constructs included PK7FWG2::*ΔNCrGEF1*, PK7FWG2::*ΔCCrGEF1*, PK7FWG2::*ΔNCrGAP1*, PK7FWG2::ΔCCrGAP1, PK7FWG2::*ΔNCrGDI2*, PK7FWG2::*ΔCCrGDI2*, or PK7FWG2 (control). Overnight Agrobacterium cultures were pelleted by centrifugation and resuspended in infiltration buffer (10 mM MES, pH 5.6, 10 mM MgCl_2_, and 200 μM acetosyringone) to a final OD₆₀₀ of 0.5–0.6. The suspension was incubated at 28°C for 4 hours before infiltration. To enhance transient expression, *Agrobacterium* cultures containing the p19 RNA silencing suppressor construct were mixed at a 1:1 ratio with Agrobacterium cultures containing the overexpression or control constructs. Infiltration was performed on the first pair of leaves using vacuum infiltration, with the leaf underside gently pinched using a needle to facilitate infiltration. Following infiltration, the plants were covered and maintained in the dark for 48 hours. Leaf samples from the first and second pairs were harvested 72 hours post-infiltration and immediately stored at -80 °C for further analysis.

### Gene Expression Measurements

For tissue specific expression analysis, total RNA from leaves, roots, stems, siliques, flower buds and flower were extracted using Trizol reagent (Sigma Aldrich, USA) following the manufacturer’s instructions. Similarly, for determining the expression of genes in MeJA elicitation, VIGS, and overexpression experiments, leaves were collected and used for total RNA extraction. cDNA was synthesized using High-capacity cDNA Reverse transcription kit (Applied Biosystems, USA) following manufacturer’s instruction, and qRT-PCR was carried out as described previously (6). The primers used for quantification of *CrGEF1, CrGAP1 and CrGDI2* transcripts were designed outside of the gene region used for cloning into pTRV2. The experimentally validated reference gene *CrN2227* was used as endogenous control. The PCR conditions were as follows: 94°C for 10 min for one cycle, followed by 40 cycles of 94°C for 15 sec, and 60°C for 1 min. Fold-change differences in gene expression were analysed using the comparative cycle threshold (C_t_) method. Relative quantification was carried out by calculating Ct to determine the fold difference in gene expression [ΔC_t_ target – ΔC_t_ calibrator]. The relative level was determined as 2_ΔΔCT.

### Metabolite Analyses and Quantification

For VIGS experiments, metabolites were extracted from the first pair of leaves of plants at the four-to-six-leaf stage (21 days post-inoculation (dpi) of two-leaf stage seedlings). In transient overexpression experiments, metabolites were extracted from the fully developed first pair of leaves of 1-month-old plants. All metabolite levels in experimental samples were compared against those of corresponding control samples at the same developmental stage. The quantification of secologanin, vindoline, catharanthine, ajmalicine, vindolinine, epi-vindoline, and ajmalicine was performed using high-performance liquid chromatography (HPLC) equipped with a photodiode array (PDA) detector. A Shimadzu HPLC system (Model: SCL-10AVP, Japan) with a C_18_ symmetry reverse-phase column (5 μm, 250 mm × 4.6 mm; Waters, Milford, MA, USA) was used for all analyses. Secologanin was extracted and quantified following Tikhomiroff and Jolicoeur (43) with minor modifications. Fresh leaf tissue (50 mg) was grounded in 1.5 ml of methanol, sonicated for 60 minutes, and centrifuged. The supernatant was decolorized with activated charcoal, transferred to a fresh tube, and evaporated to dryness. The dried residue was dissolved in 20μl of methanol and analyzed via HPLC. The mobile phase consisted of a 15:85 (v/v) mixture of gradient-grade acetonitrile and 0.1 M phosphoric acid (pH 2.0), delivered at a flow rate of 1.5 ml/min in isocratic mode for 20 minutes. Data were recorded at 238 nm. Vindoline, catharanthine, vindolinine, epi-Vindoline, and ajmalicine were extracted based on the protocol of (44), with minor modifications. Oven-dried leaf tissue (10 mg) was grounded in 2 ml of methanol and incubated at 55°C for 2 hours with intermittent shaking. The extract was filtered and evaporated to dryness, and the residue was dissolved in 700 μl of 2.5% H₂SO₄. The sample underwent two rounds of extraction with equal volumes of ethyl acetate, retaining the aqueous phase each time. The pH of the aqueous phase was then adjusted to 9.0 using NH₄OH, followed by alkaloid extraction with ethyl acetate. The organic phase was evaporated to dryness, and the alkaloid-containing residue was dissolved in 20 μl of methanol for HPLC analysis. The HPLC mobile phase included solvent A (0.1 M ammonium acetate buffer, pH 7.3) and solvent B (gradient-grade acetonitrile). The gradient program began with a 70:30 ratio of A at a flow rate of 1 ml/min for 5 minutes, ramped linearly to 36:64 at 1.4 ml/min over the next 5 minutes, adjusted to 20:80 at 1.4 ml/min for another 5 minutes, and returned to 70:30 at 1 ml/min for the final 5 minutes. Data were recorded at 254 nm.

### Statistical Analysis

Average means, standard error (SE) and number of replicates (biological and technical) for each experiment was used to determine the statistical significance using GraphPad9 Prism software. The statistical significance of differences between control and treated samples were tested using one-way and two-way ANOVA with Sidak test.

## Acknowledgments

This work was partly supported by MLP0003 project. A.S. is a recipient of a research fellowship from University Grants Commission (UGC). The authors are thankful to Dr. Ramu Vemana for sparing pDEST22 and pDEST32 vectors. Authors are also thankful to Dr. Sumit Ghosh for sparing pGWB441 vector, and Rajendra Patel for help with confocal analysis. The authors also express their sincere gratitude to the Director, CSIR-CIMAP for support throughout the study. Institutional communication number for this article is CIMAP/Pub/2024/167. Authors declare no conflict of interest.

## Author Contributions

DAN planned and designed the research. AS, SM, PG and DP performed experiments. AS and DAN analyzed and interpreted the data and wrote the manuscript.

## Supporting Information

**Figure S1.**
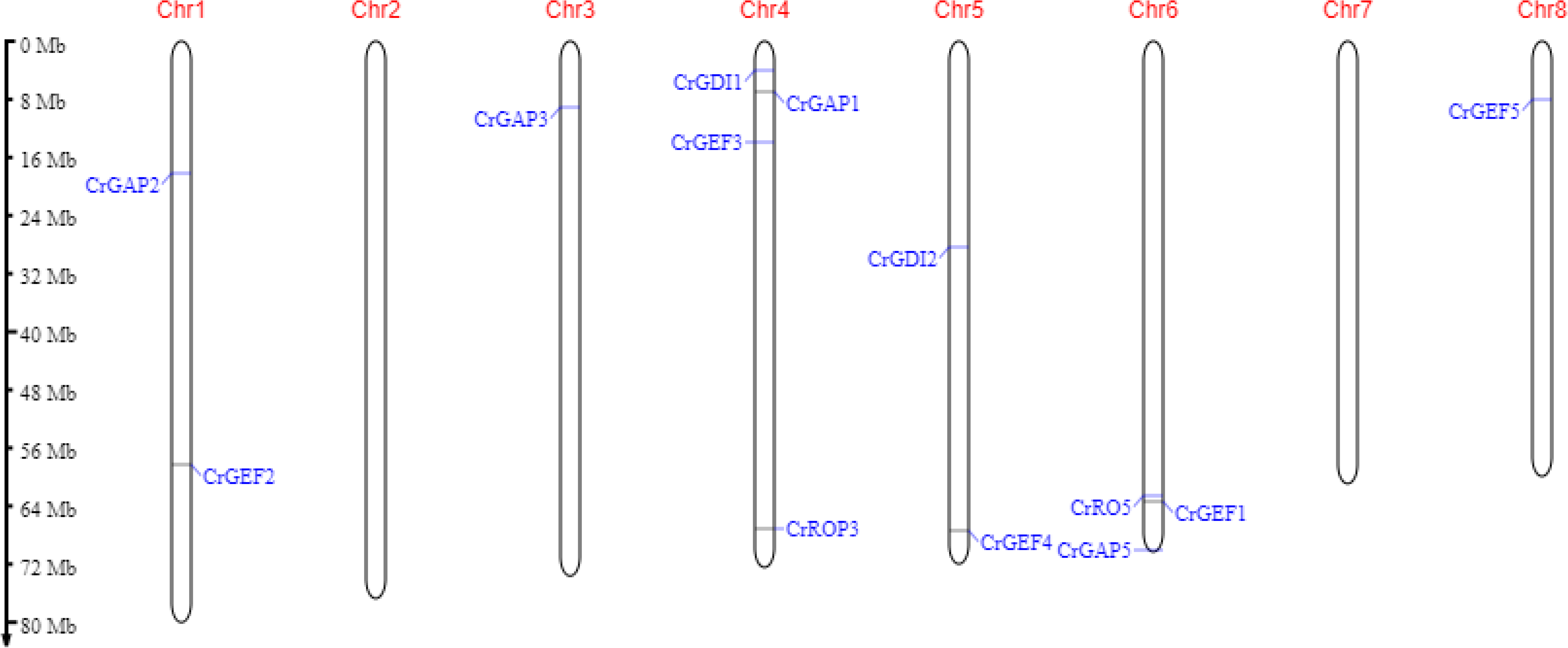
Diagram showing the location of *CrROP3/5, CrGEFs, CrGAPs* and *CrGDIs* in *Catharanthus roseus* genome as obtained from MapGene2Chrom v2 (http://mg2c.iask.in/mg2c_v2.1/index.html)

**Figure S2.**
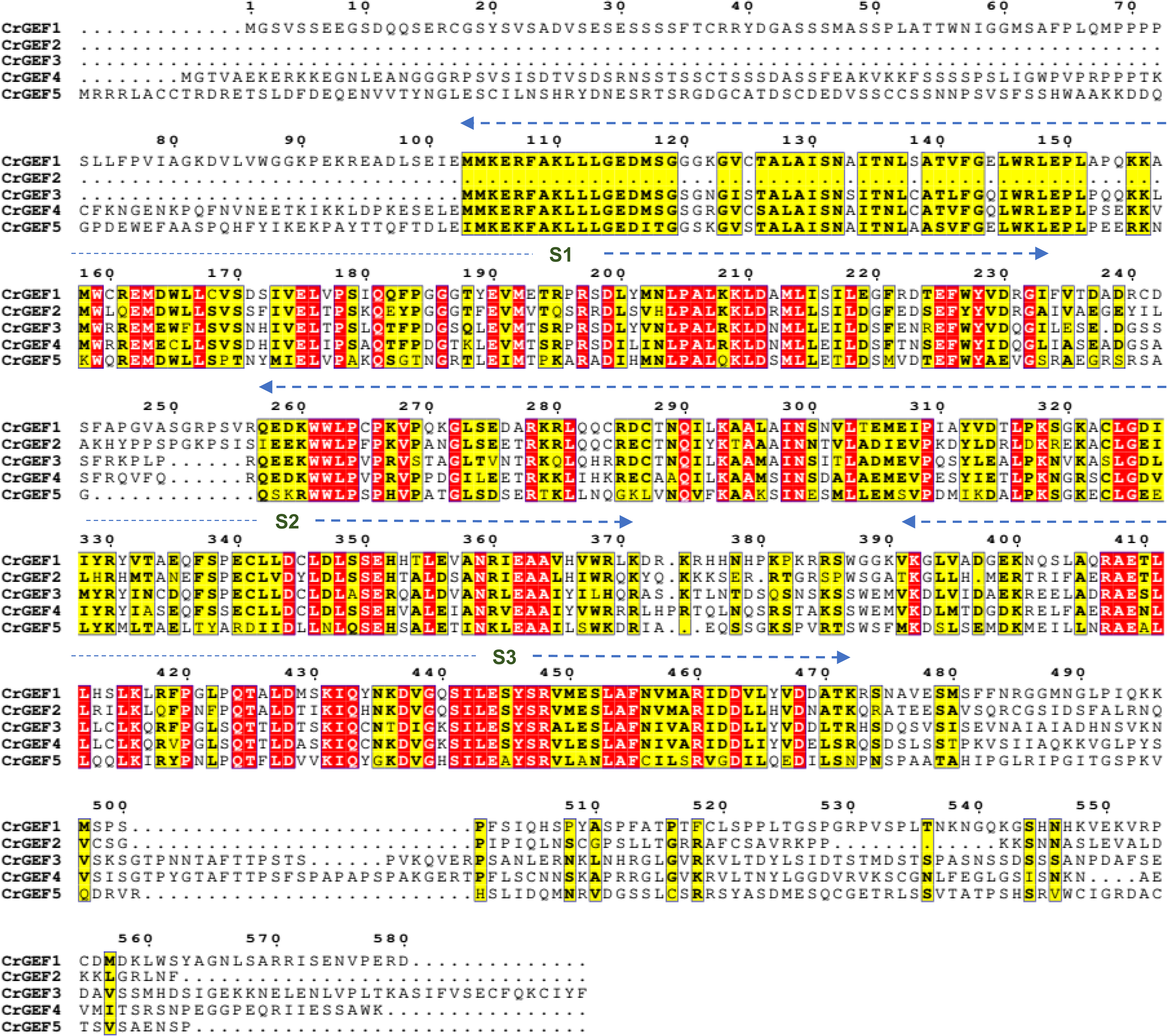
Multiple sequence alignment of CrGEFs was performed using Clustal W (Thompson et al. 1994) with default parameters through https://www.genome.jp/tools-bin/clustalw. Flashy coloring was done with ESPript 3.0 (https://espript.ibcp.fr/ESPript/ESPript/) in which amino acids in seprate boxes indicate identical and conserved residues in all the five sequences. Dotted lines above aligned sequences indicate highly conserved regions in a PRONE domain composed of three subdomains (S1, S2, and S3).

**Figure S3.**
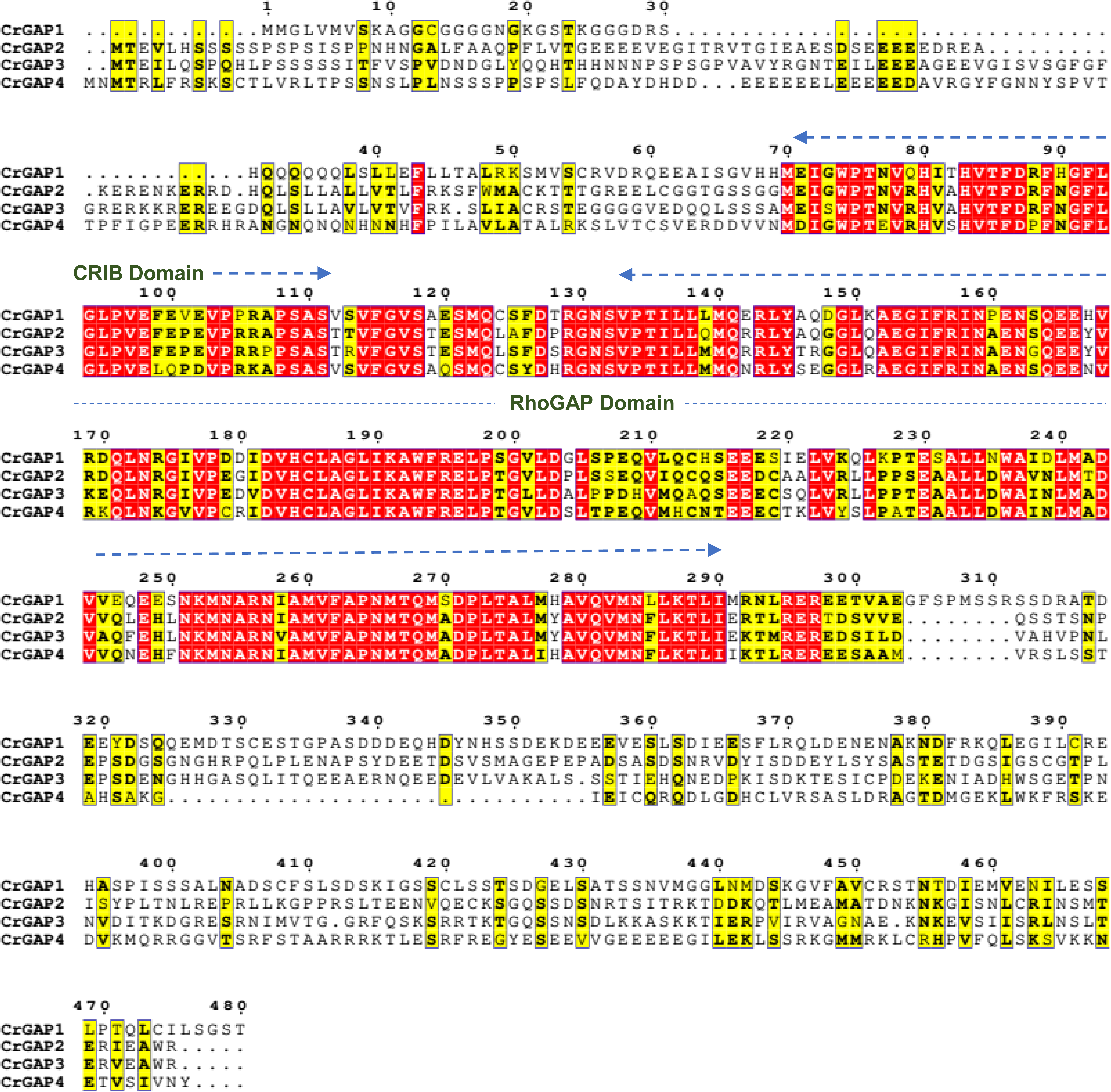
Multiple sequence alignment of CrGAPs was performed using Clustal W (Thompson et al. 1994) with default parameters through https://www.genome.jp/tools-bin/clustalw. Flashy coloring was done with ESPript 3.0 (https://espript.ibcp.fr/ESPript/ESPript/) in which amino acids in seprate boxes indicate identical and conserved residues in all the four sequences. Dotted lines above aligned sequences indicate highly conserved regions CRIB and RhoGAP domain.

**Figure S4.**
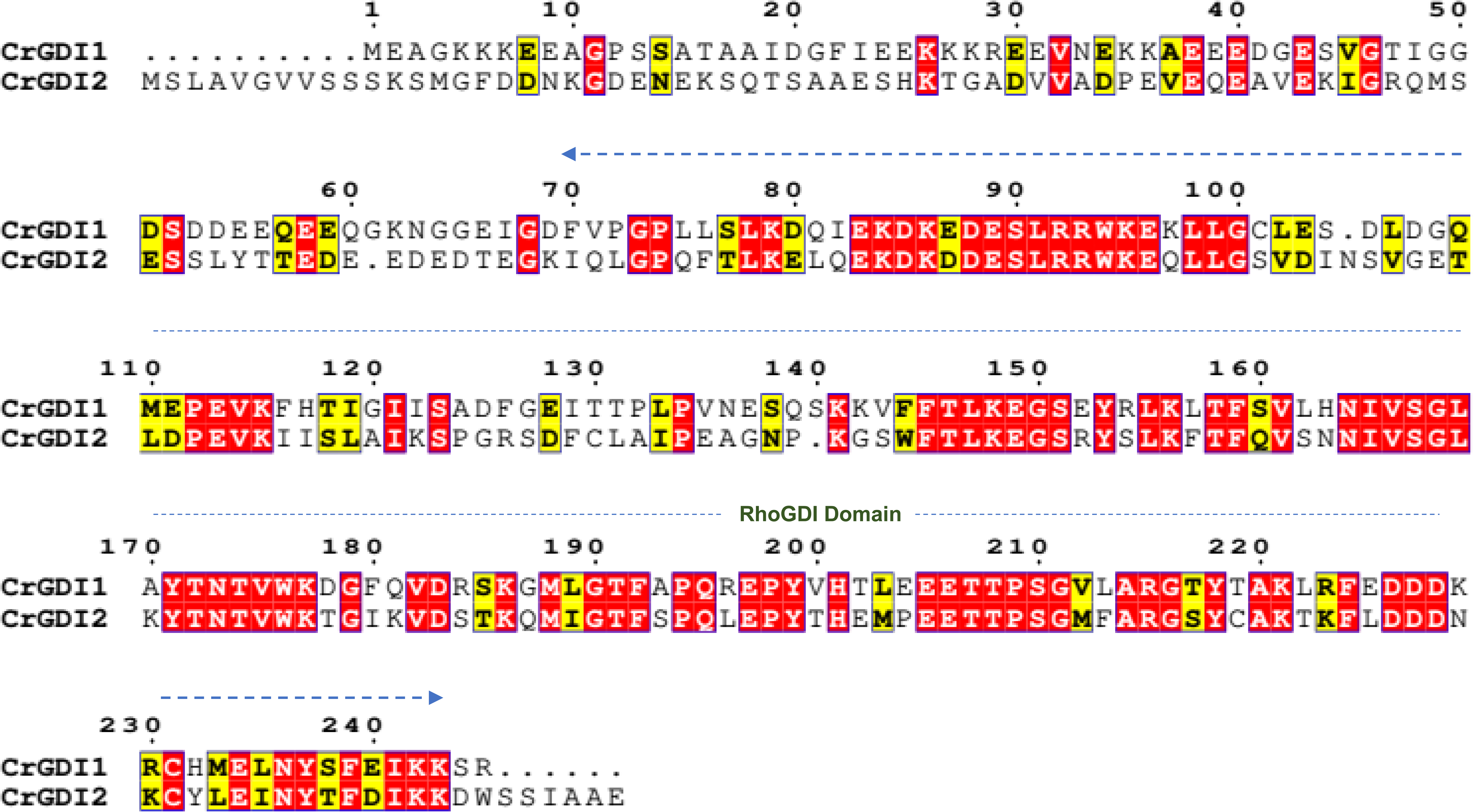
Multiple sequence alignment of CrGDIs was performed using Clustal W (Thompson et al. 1994) with default parameters through https://www.genome.jp/tools-bin/clustalw. Flashy coloring was done with ESPript 3.0 (https://espript.ibcp.fr/ESPript/ESPript/) in which amino acids in seprate boxes indicate identical and conserved residues in all the two sequences. Dotted lines above aligned sequences indicate highly conserved regions RhoGDI domain.

**Figure S5.**
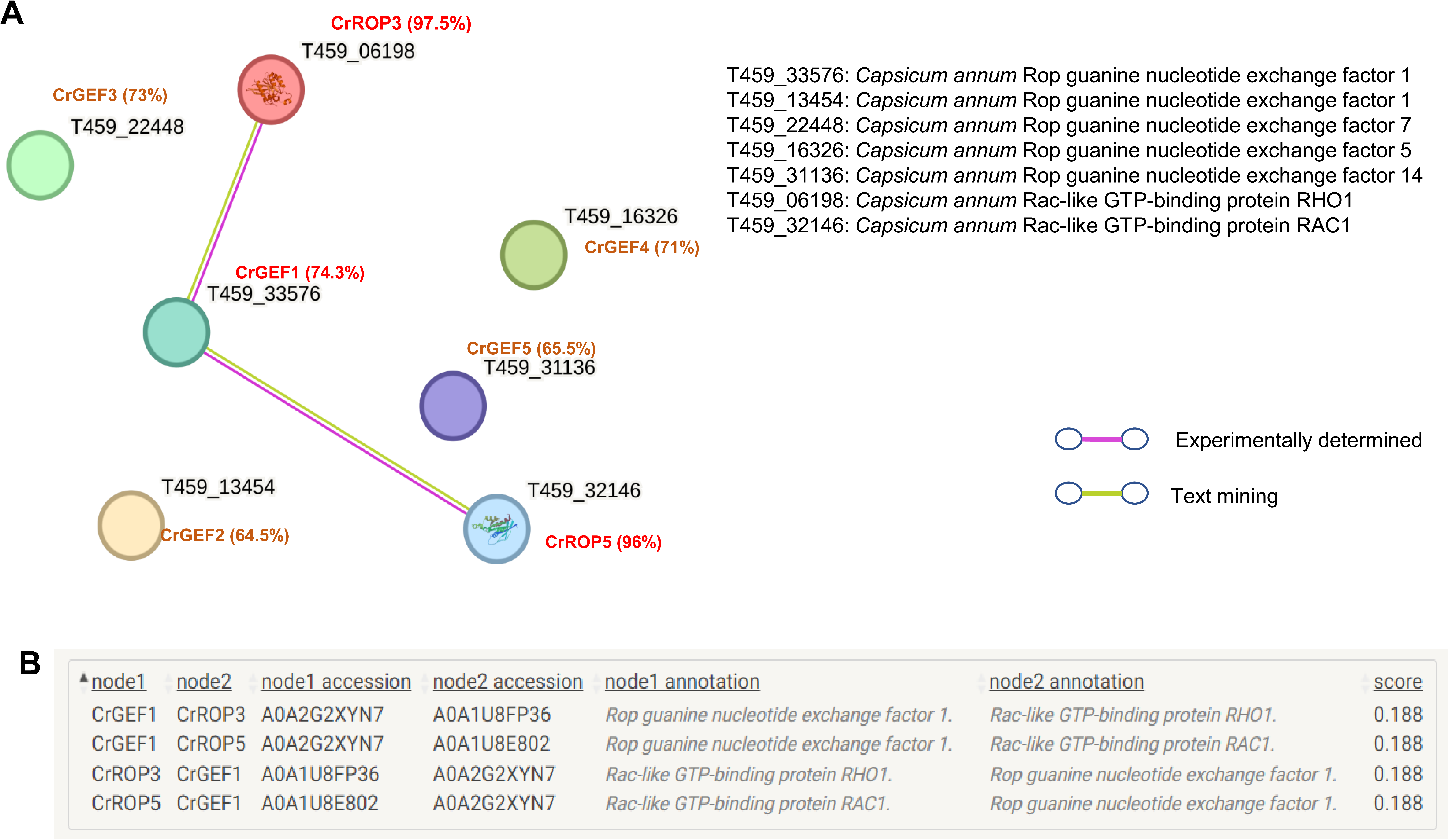
*In silico* analysis of protein-protein interaction between CrROP3 or CrROP5 and CrGEF1 or CrGEF2 or CrGEF3 or CrGEF4 or CrGEF5 using online prediction software STRING 12.0. tool (https://string-db.org/). (A) Green and purple lines indicate possible interaction of CrGEF1 (homolog of *Capsicum annum* RopGEF1) with CrROP3 (homolog of *C. annum* RHO1) or CrROP5 (homolog of *C. annum* RAC1). (B) Score for *in silico* interaction.

**Figure S6.**
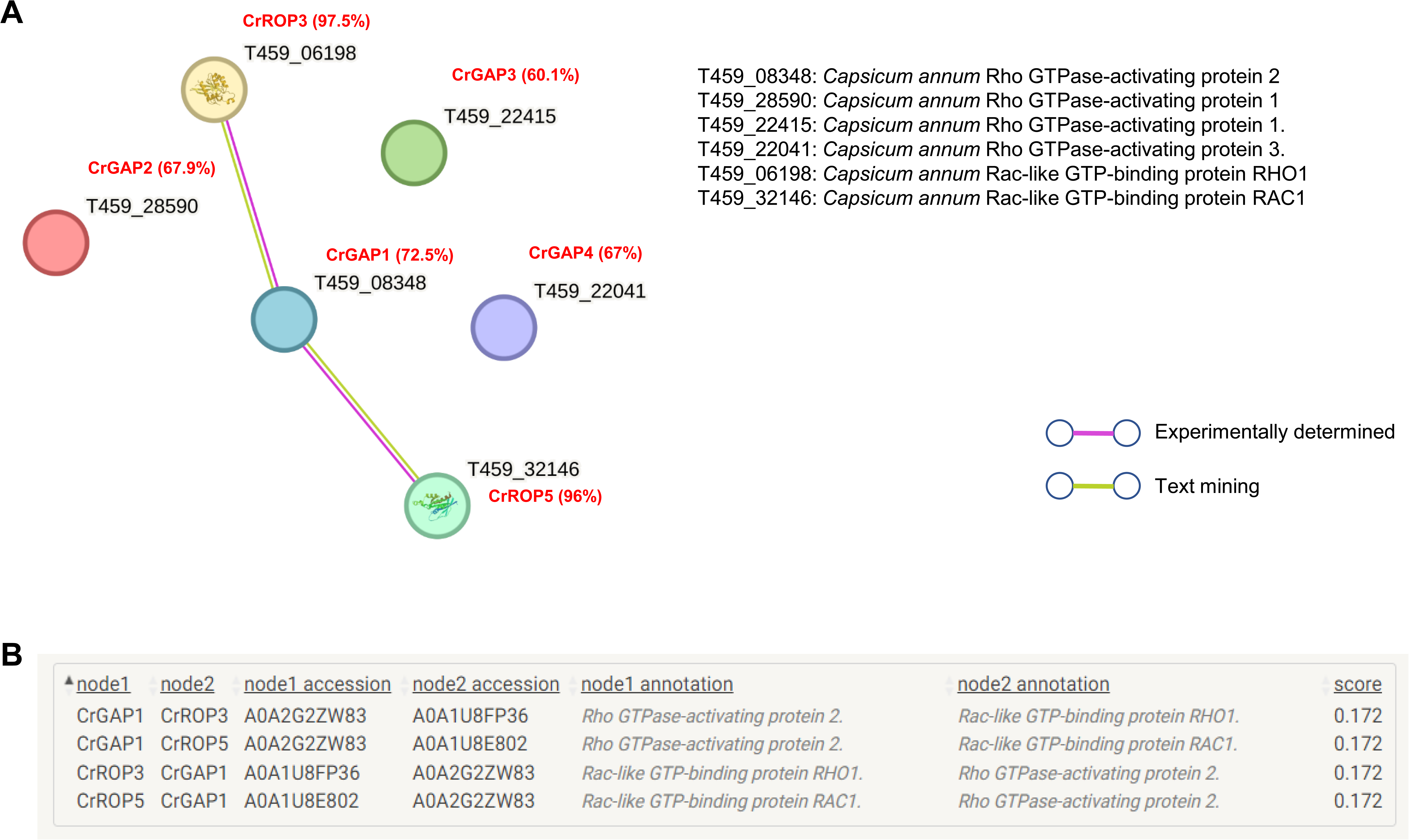
*In silico* analysis of protein-protein interaction between CrROP3 or CrROP5 and CrGAP1 or CrGAP2 or CrGAP3 or CrGAP4 using online prediction software STRING 12.0. tool (https://string-db.org/). (A) Green and purple lines indicate possible interaction of CrGAP1 (homolog of *Capsicum annum* RhoGAP2) with CrROP3 (homolog of *C. annum* RHO1) or CrROP5 (homolog of *C. annum* RAC1). (B) Score for *in silico* interaction.

**Figure S7.**
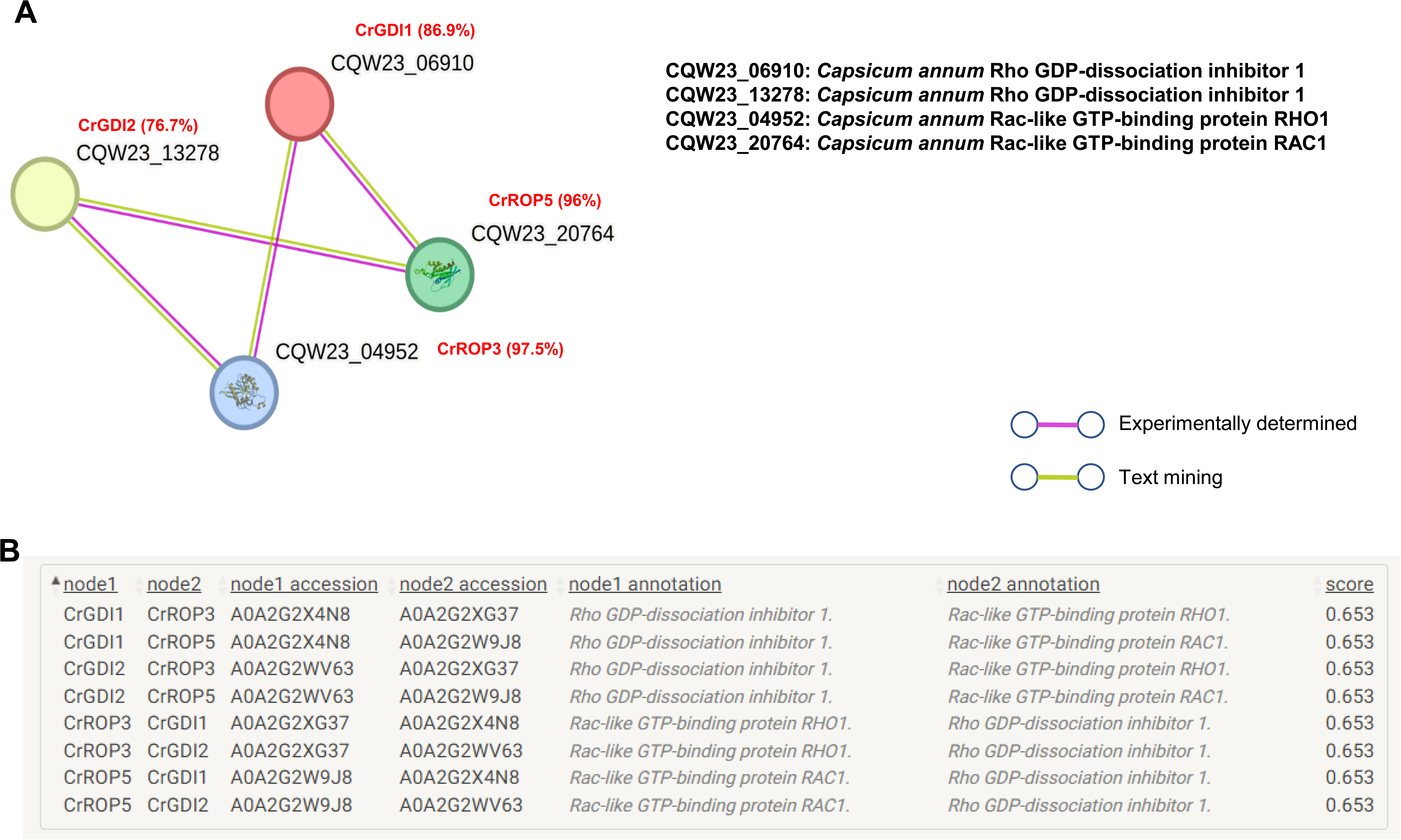
*In silico* analysis of protein-protein interaction between CrROP3 or CrROP5 and CrGDI1 or CrGDI2 using online prediction software STRING 12.0. tool (https://string-db.org/). (A) Green and purple lines indicate possible interaction of CrGDI1 (homolog of *Capsicum annum* RhoGDI1) or CrGDI2 (homolog of *Capsicum annum* RhoGDI1) with CrROP3 (homolog of *C. annum* RHO1) or CrROP5 (homolog of *C. annum* RAC1). (B) Score for *in silico* interaction.

**Figure S8.**
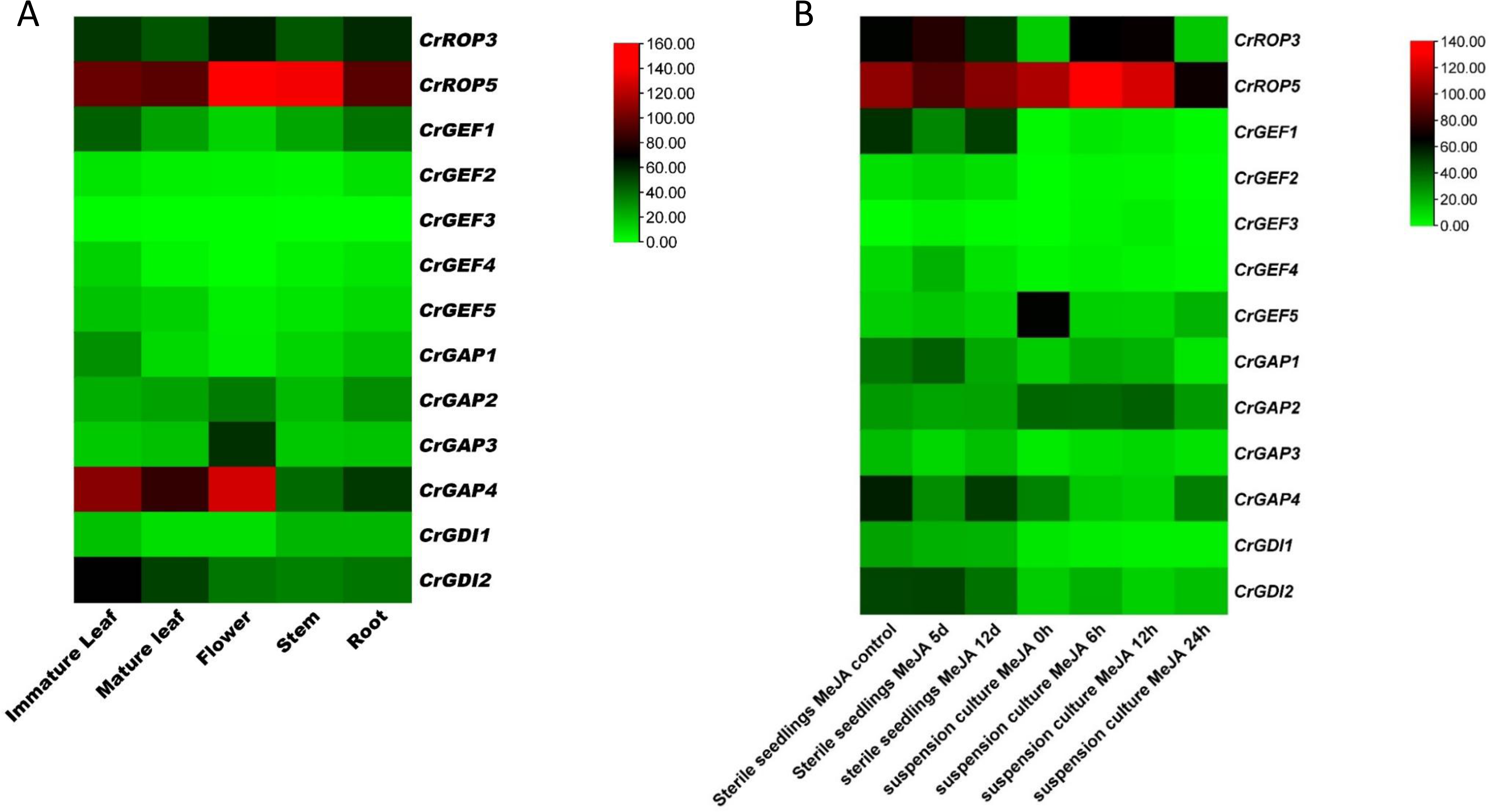
Heat map (generated using the software TBtools) showing *in silico* expression profile of *CrROP3/5, CrGEF1-5, CrGAP1-5* and *CrGDI1-2* in different tissues (A), and in response to methyl jasmonate (MeJA) in seedlings and suspension culture (B). The heat map is representative of the log_2_ values of FPKM which were obtained from MPGR database (http://mpgr.uga.edu/).

**Figure S9.**
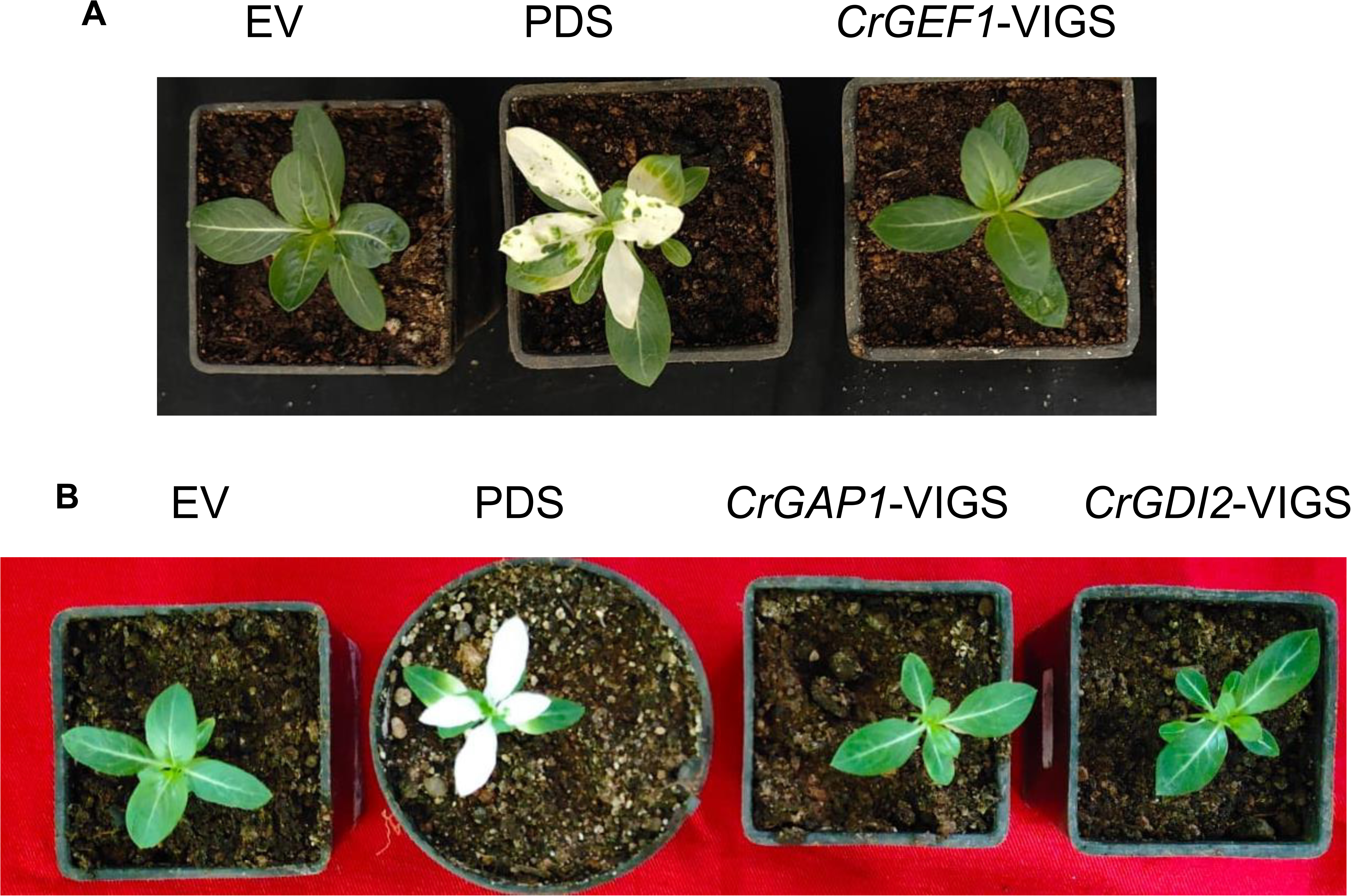
Tobacco rattle virus-mediated silencing in *C. roseus*. (A) Representative *C. roseus* plants 3 weeks post-infiltration with 1:1 of pTRV1/pTRV2 (empty vector, EV), pTRV1/pTRV2-CrPDS (CrPDS-vigs) and pTRV1/pTRV2-CrGEF1 (CrGEF1-vigs). (B) Representative C. roseus plants 3 weeks post-infiltration with 1:1 of and pTRV1/pTRV2 (empty vector, EV),. pTRV1/pTRV2-CrPDS (CrPDS-vigs), pTRV1/pTRV2-CrGAP1 (CrGAP1-vigs) and pTRV1/Ptrv2-CrGDI2 (CrGDI2-vigs).

**Figure S10.**
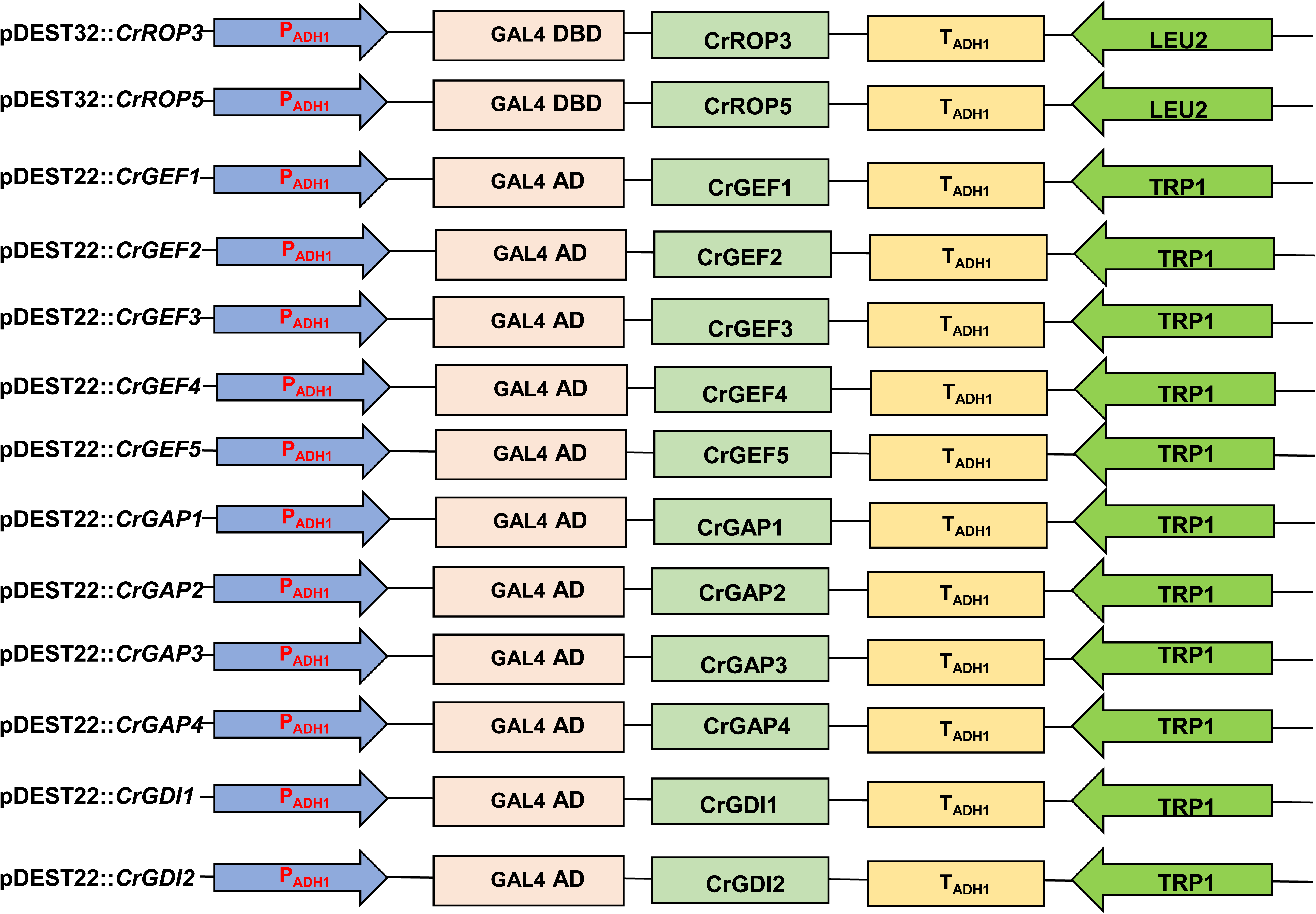
Schematic diagrams of Yeast two-hybrid assay constructs generated and used in this study.

**Figure S11.**
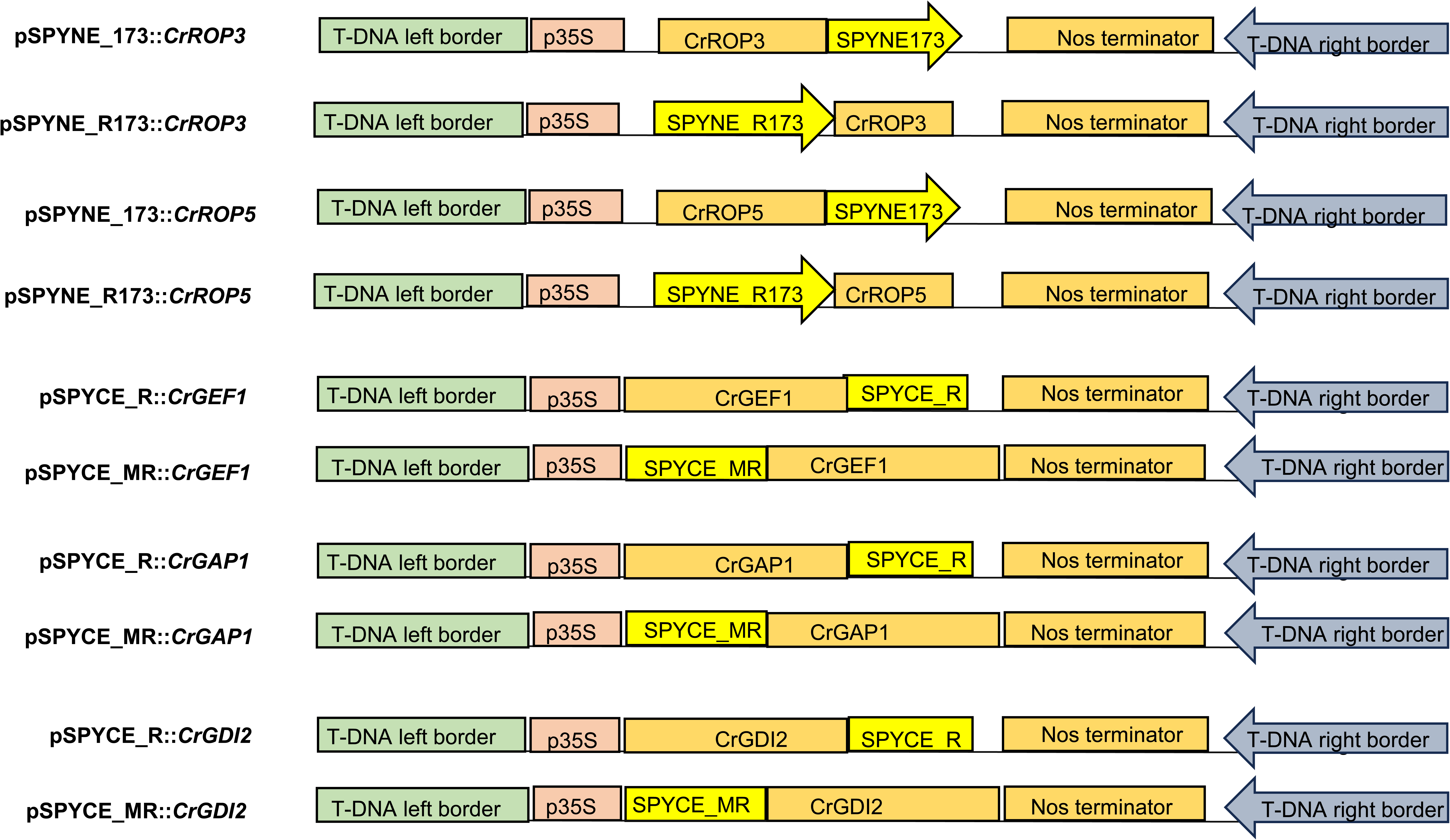
Schematic diagrams of Bimolecular fluorescence complementation (BiFC) assay constructs generated and used in this study.

**Figure. S12.**
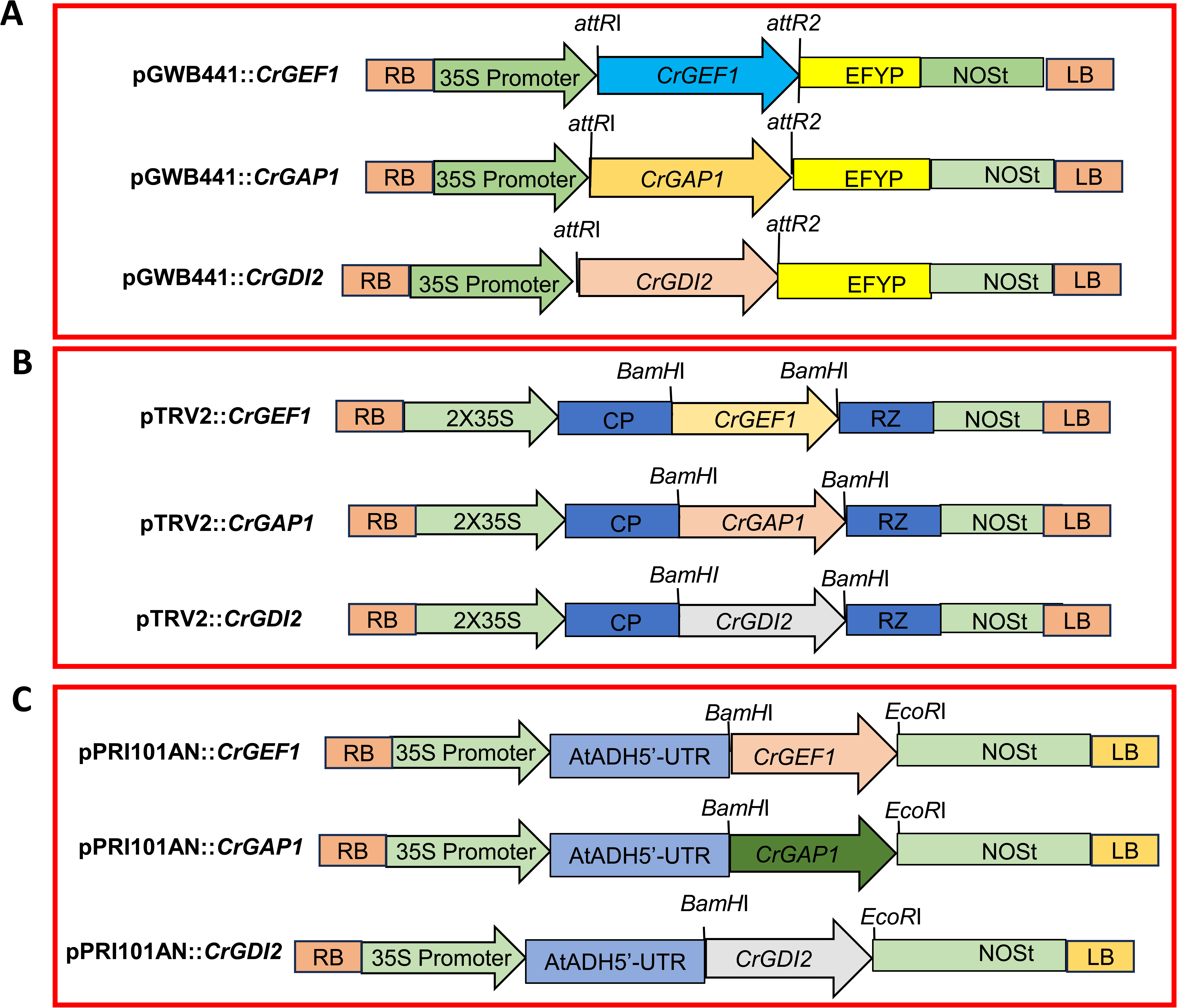
Schematic diagrams of subcellular localization, virus induce gene silencing and overexpression constructs generated and used in this study. (A) pGWB441-derived constructs used for subcellular localization. (B) TRV2-derived VIGS constructs. (C) pRI101AN-derived plant overexpression constructs. NOST, Nopaline synthase terminator; UTR, untranslated region; CP, coat protein; RZ, self-cleaving ribozyme RB, Right Border; LB, Left border.

**Figure S13.**
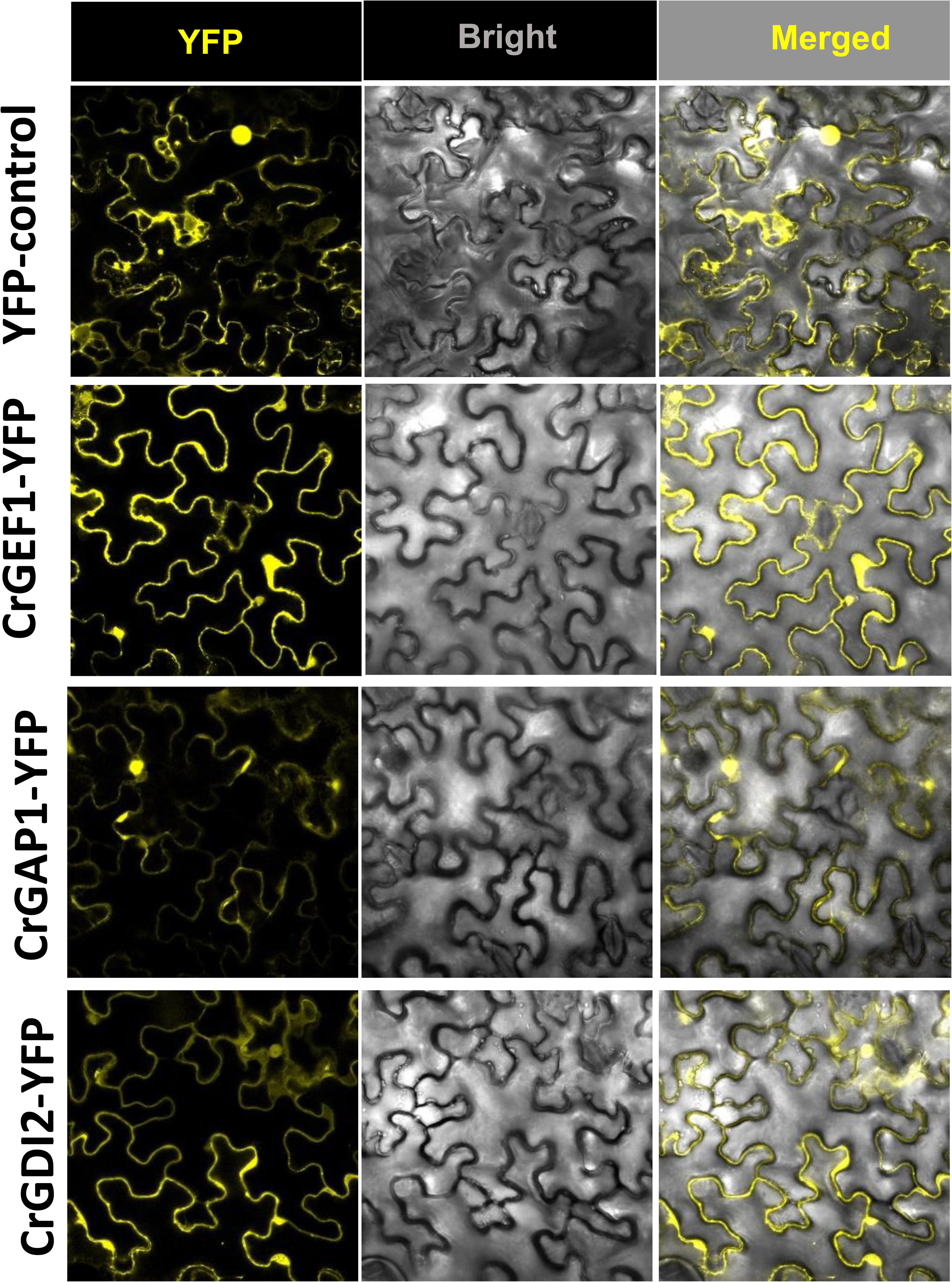
Additional confocal images of CrGEF1-YFP, CrGAP1-YFP and CrGDI2-YFP expressing cells.

**Supplemental Table 1.**
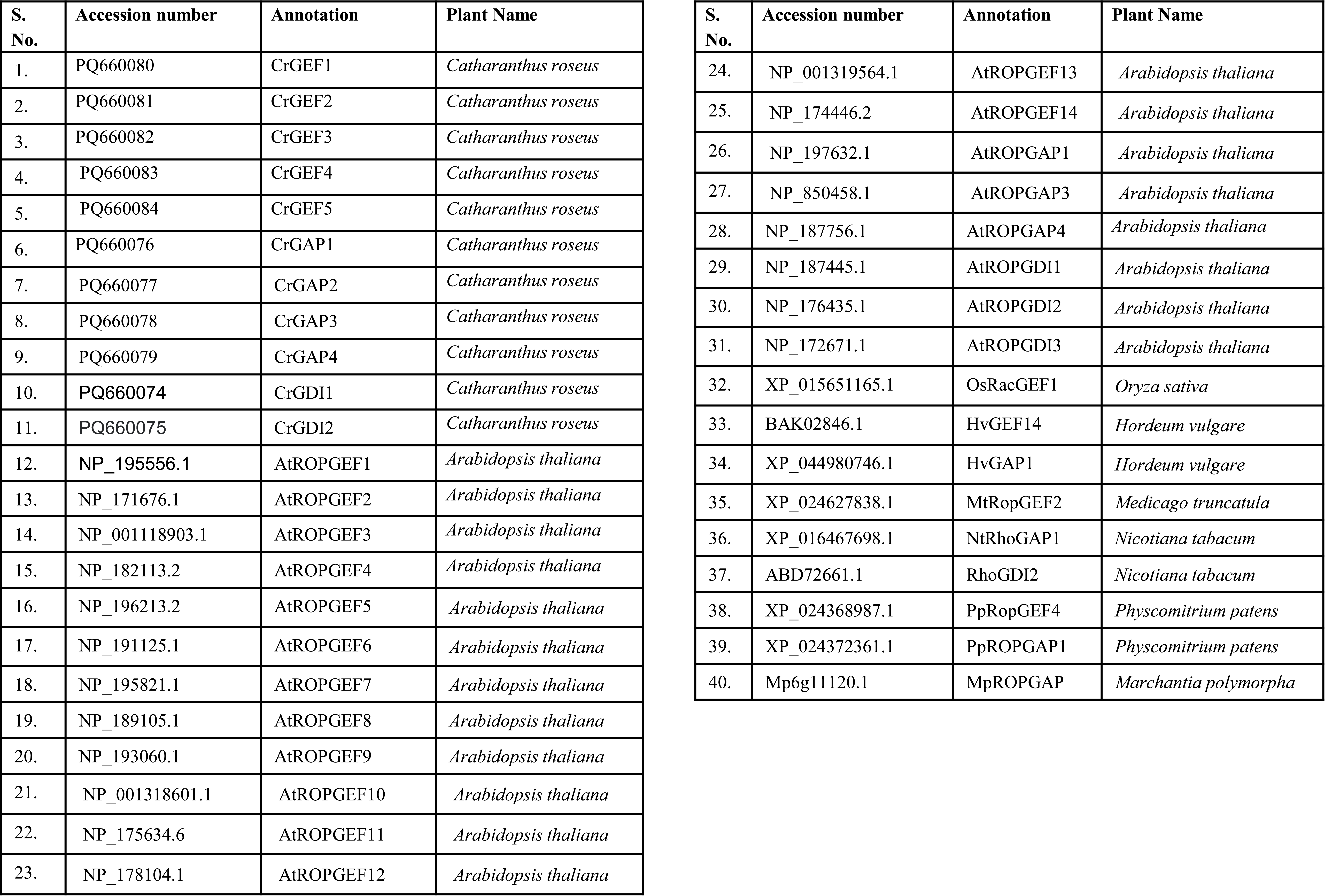
Accession numbers and gene annotations of the sequences used in phylogenetic analysis.

**Supplemental Table 2.**
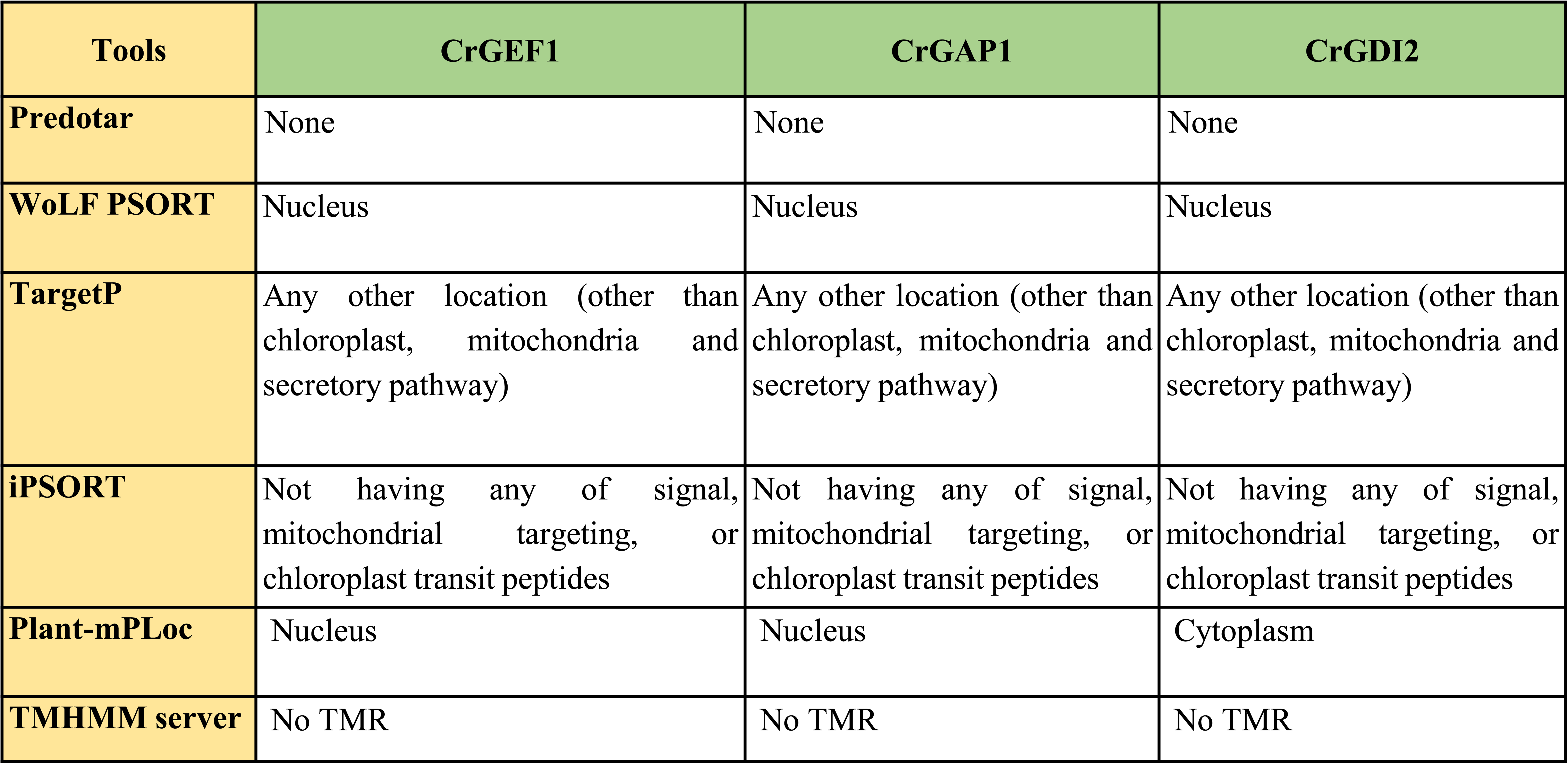
*In silico* subcellular localization prediction of CrGEF1, CrGAP1, and CrGDI2 using different software tools.

**Supplemental Table 3.**
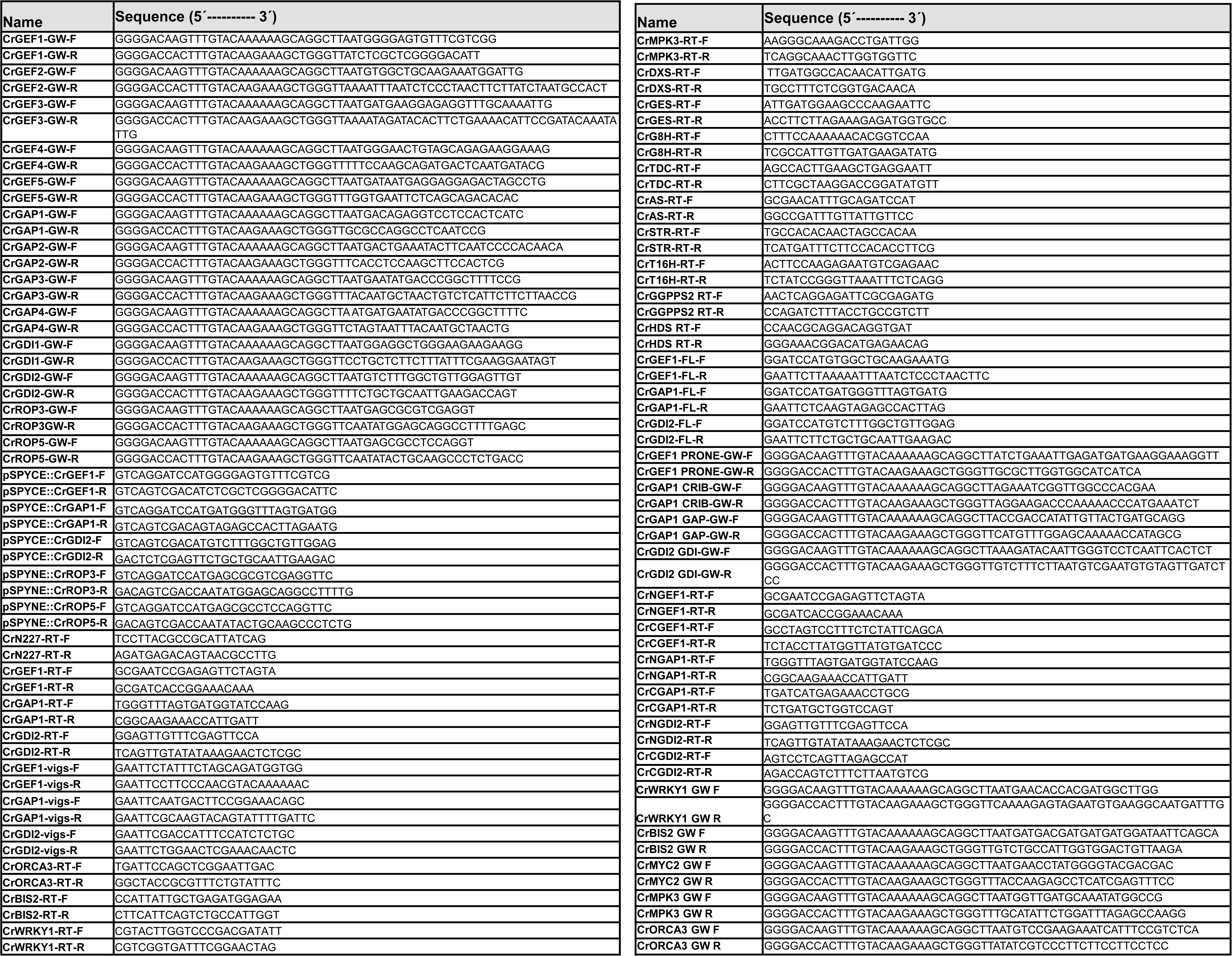
List of oligonucleotide primers used in this study.

## Notes

### Competing Interest Statement

The authors have declared no competing interest.

